# Parallel reconstruction of the excitatory and inhibitory inputs received by single neurons reveals the synaptic basis of recurrent spiking

**DOI:** 10.1101/2023.01.06.523018

**Authors:** Julian Bartram, Felix Franke, Sreedhar Saseendran Kumar, Alessio Paolo Buccino, Xiaohan Xue, Tobias Gänswein, Manuel Schröter, Taehoon Kim, Krishna Chaitanya Kasuba, Andreas Hierlemann

## Abstract

Self-sustained recurrent activity in cortical networks is thought to be important for multiple crucial processes, including circuit development and homeostasis. Yet, the precise relationship between synaptic input patterns and the spiking output of individual neurons remains largely unresolved. Here, we developed, validated and applied a novel in vitro experimental platform and analytical procedures that provide – for individual neurons – simultaneous excitatory and inhibitory synaptic activity estimates during recurrent network activity. Our approach combines whole-network high-density microelectrode array (HD-MEA) recordings from rat neuronal cultures with patch clamping and enables a comprehensive mapping and characterization of active incoming connections to single postsynaptic neurons. We found that, during network states with excitation(E)-inhibition(I) balance, postsynaptic spiking often coincided with the maxima of fast fluctuations in the input E/I ratio. These spike-associated E/I ratio escalations were largely due to a rapid bidirectional change in synaptic inhibition that was modulated by the network-activity level. Our approach also uncovered the underlying circuit architecture and we show that individual neurons received a few key inhibitory connections – often from special hub neurons – that were instrumental in controlling postsynaptic spiking. Balanced network theory predicts dynamical regimes governed by small and rapid input fluctuation and featuring a fast neuronal responsiveness. Our findings – obtained in self-organized neuronal cultures – suggest that the emergence of these favorable regimes and associated network architectures is an inherent property of cortical networks in general.

## Introduction

Neurons typically receive a continuous bombardment by orchestrated excitatory and inhibitory synaptic inputs, which ultimately determines postsynaptic spiking. Such a continuous input activation is a basic operational principle of cortical networks that has been observed in awake animals (Destexhe et al. 2003; Steriade et al. 2001; Timofeev et al. 2001), up states recorded during slow-wave sleep (Steriade et al. 1993, 2001) and in brain slices (Haider and McCormick 2009; Sanchez-Vives and McCormick 2000; Shu et al. 2003), and periods of heightened network activity in cell cultures (Liu 2004; Van Pelt et al. 2004; Wagenaar et al. 2006a,b; Xue et al. 2022). The observed input barrages can be generated – to a large extent or even completely in the in vitro cases – by spontaneous recurrent network activity. Self-maintained network activity is thought to be important for such diverse processes as synaptic homeostasis (Bartram et al. 2017), a mediation of circuit refinements (Pan and Monje 2020) and working memory (Seamans et al. 2003), but the circuit properties and dynamics that give rise to and shape spontaneous activity are poorly understood in biological neural networks.

Numerous studies have shown that the excitatory (E) and inhibitory (I) inputs received by individual cortical neurons are approximately balanced through local circuit interactions during both evoked and spontaneous network activity (Haider and McCormick 2009; Isaacson and Scanziani 2011; Liu 2004; Shu et al. 2003). The approximate input balance raises the question of precisely which synaptic activation patterns are actually associated with postsynaptic spiking. The answer to this question would provide insights into the dynamical regimes in which networks operate and has, therefore, important implications for network function (Ahmadian and Miller 2021). Theoretical work suggests that balanced networks can potentially assume multiple different dynamical states (Brunel 2000). When global spiking is asynchronous and neuronal firing is irregular – as often observed in cortical networks during wakefulness – neural networks may operate in a so-called fluctuation-driven regime. In this dynamical state, the mean neuronal membrane potential is just below the action potential threshold and fast input fluctuations yield brief threshold crossings (Ahmadian and Miller 2021; Amit and Brunel 1997; Brunel 2000; Van Vreeswijk and Sompolinsky 1996). Similarly, in regimes with increased input synchronization and oscillatory dynamics (e.g. in the gamma-frequency range), the neuronal membrane potential is predominantly near the action potential threshold (Buzsáki and Wang 2012; Hasenstaub et al. 2005). The described subthreshold regimes have favorable properties: Neurons can respond rapidly and precisely to small and brief input changes, and multiple different combinations of E-I conductance changes can control spiking. Contrasting these spiking mechanisms, action potential firing with relatively short and regular inter-spike intervals in response to a prolonged suprathreshold depolarization has been described (Petersen and Berg 2016; Renart et al. 2007). During such mean-driven spiking, the spike timing is controlled by after-hyperpolarization characteristics rather than rapid input changes.

To better understand and characterize the spiking regimes that are implemented in biological neural networks, our goal was to experimentally identify the synaptic events that determine postsynaptic spiking during spontaneous network activity. For such an investigation, information on *i*) the excitatory and *ii*) inhibitory synaptic input conductances in addition to *iii*) the postsynaptic spike times is needed. However, with existing techniques, it is impractical to perform these three measurements in parallel. While the dynamic clamp technique and computational modelling may be used to investigate the modulation of spike timing by artificial synaptic conductances (Hasenstaub et al. 2005; Piwkowska et al. 2008), these approaches are limited in their ability to recapitulate the full complexity of the activities generated by biological neural networks.

To address the existing methodological shortcomings, we developed a novel experimental approach and analytical tools that provide – in parallel – a reconstruction of the excitatory and inhibitory synaptic input activity during a period of recorded postsynaptic spiking (**Fig. 1**). The result of this reconstruction is a detailed input-output characterization of individual neurons during multiple hours of recurrent network activity. We combined high-density microelectrode array (HD-MEA) recordings with patch clamping and used a cortical cell culture model that generated a rich repertoire of E-I balanced spontaneous recurrent network activity with similarities to cortical up-down state oscillations of deep sleep (Barral and Reyes 2016; Haider and McCormick 2009; Liu 2004). This in vitro model, by nature, was free of external inputs, permitted control over the network size, and, crucially, allowed for recording of spiking activity from virtually every neuron in the network. We validated this reconstruction approach and, subsequently, applied it to examine the synaptic basis of recurrent spiking – linking circuit activity and architecture.

**Figure 1.**
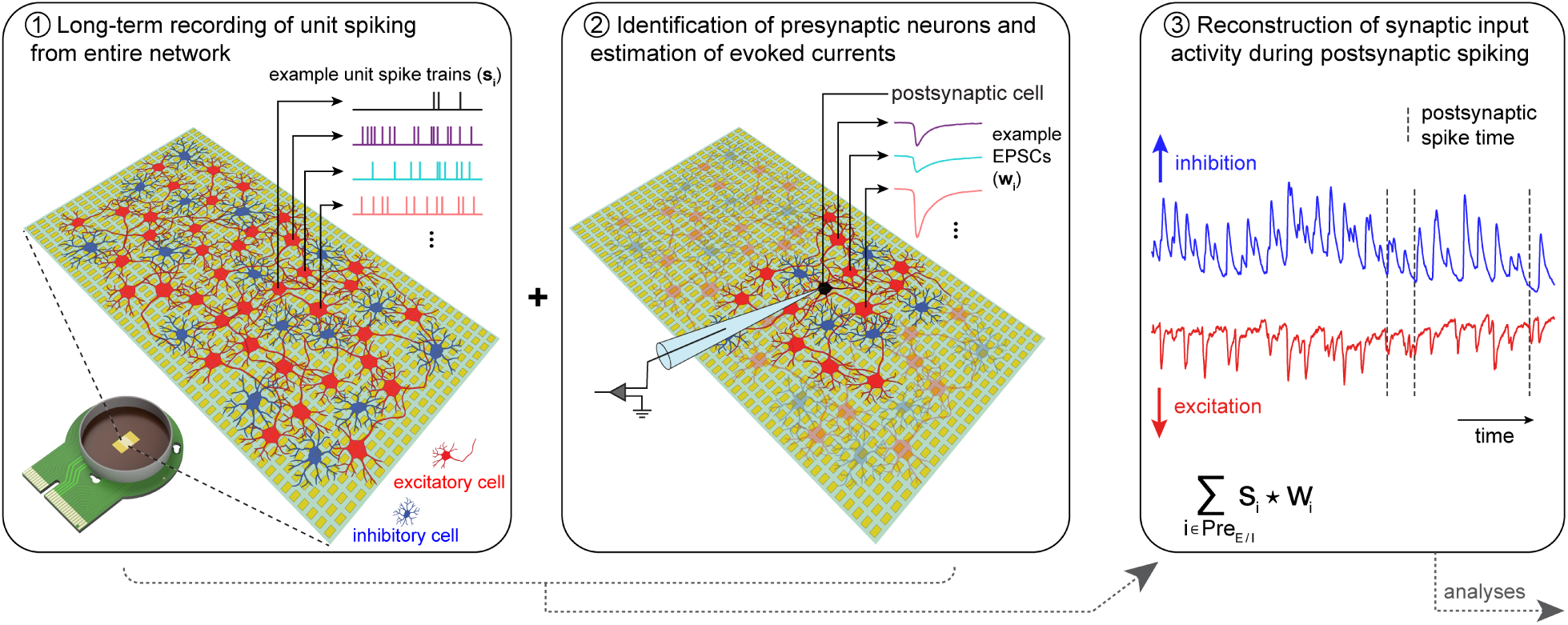
Reconstructing synaptic input activity during spontaneous network spiking. Schematic of main concept. Long-term whole-network extracellular recordings of neuronal spiking are performed using a high-density microelectrode array (HD-MEA) platform with 26’400 electrodes and 1024 channels for simultaneous readout. Next, the incoming monosynaptic connections to individual postsynaptic cells are identified using a novel regression approach applied to simultaneous HD-MEA and whole-cell patch-clamp recordings. Finally, spike trains (*s_i_*) of all identified excitatory (*Pre_E_*) or inhibitory (*Pre_I_*) presynaptic cells are convolved with their respective postsynaptic-current estimate (*w_i_*) to obtain, for several hours, parallel reconstructions of the excitatory and inhibitory synaptic input activities. Crucially, postsynaptic spike times are also available during the reconstruction period. The experimental implementation of this approach is shown in Fig. 2. Main analytical steps are introduced in Fig. 3 and 4.

## Results

### Reconstructing synaptic input activity during spontaneous network spiking

The key steps to reconstruct the excitatory and inhibitory synaptic input activity of an individual postsynaptic neuron, during a period of recorded postsynaptic spiking, are as follows (**Fig. 1**): *i)* Acquire long-term whole-network extracellular recordings of spontaneous neuronal spiking. *ii)* Identify the incoming monosynaptic connections onto individual postsynaptic cells, and calculate the mean evoked postsynaptic currents for each connection. *iii)* Identify the postsynaptic and presynaptic neurons in the long-term extracellular recording period to obtain parallel spike trains. With the spike trains of the presynaptic cells and the corresponding estimates of the evoked currents, reconstruct the excitatory and inhibitory synaptic input activity experienced by the target cell during postsynaptic spiking. In the first part of this study, we will elaborate on how each of these steps was implemented.

Our experimental pipeline is depicted in **Fig. 2**. Primary rat cortical neurons were plated on a HD-MEA chip featuring 26’400 electrodes and 1024 channels for simultaneous readout (**Fig. 2A**). Following network maturation, brief sequential recordings – covering together the entire HD-MEA chip – were performed to identify electrodes that detected neuronal activity. Active electrodes were subsequently selected for long-term recording of network-wide spiking. Extracellular data were acquired for at least 3 h (**Fig. 2B**) and spike sorted to identify individual units (**Fig. 2C**). Next, selected cells were patched, and paired HD-MEA and patch-clamp recordings were obtained (**Fig. 2D**). Postsynaptic spikes were recorded in whole-cell current-clamp or cell-attached mode, and using these spike times to generate the spike-triggered average of each HD-MEA electrode trace revealed the extracellular signature of the patched cell (here referred to as the cell or unit ‘footprint’). This footprint was matched to a unit footprint from the spike-sorted long-term recording that preceded the patch-clamp experiment in order to obtain the long-term spike train of the patched cell.

**Figure 2.**
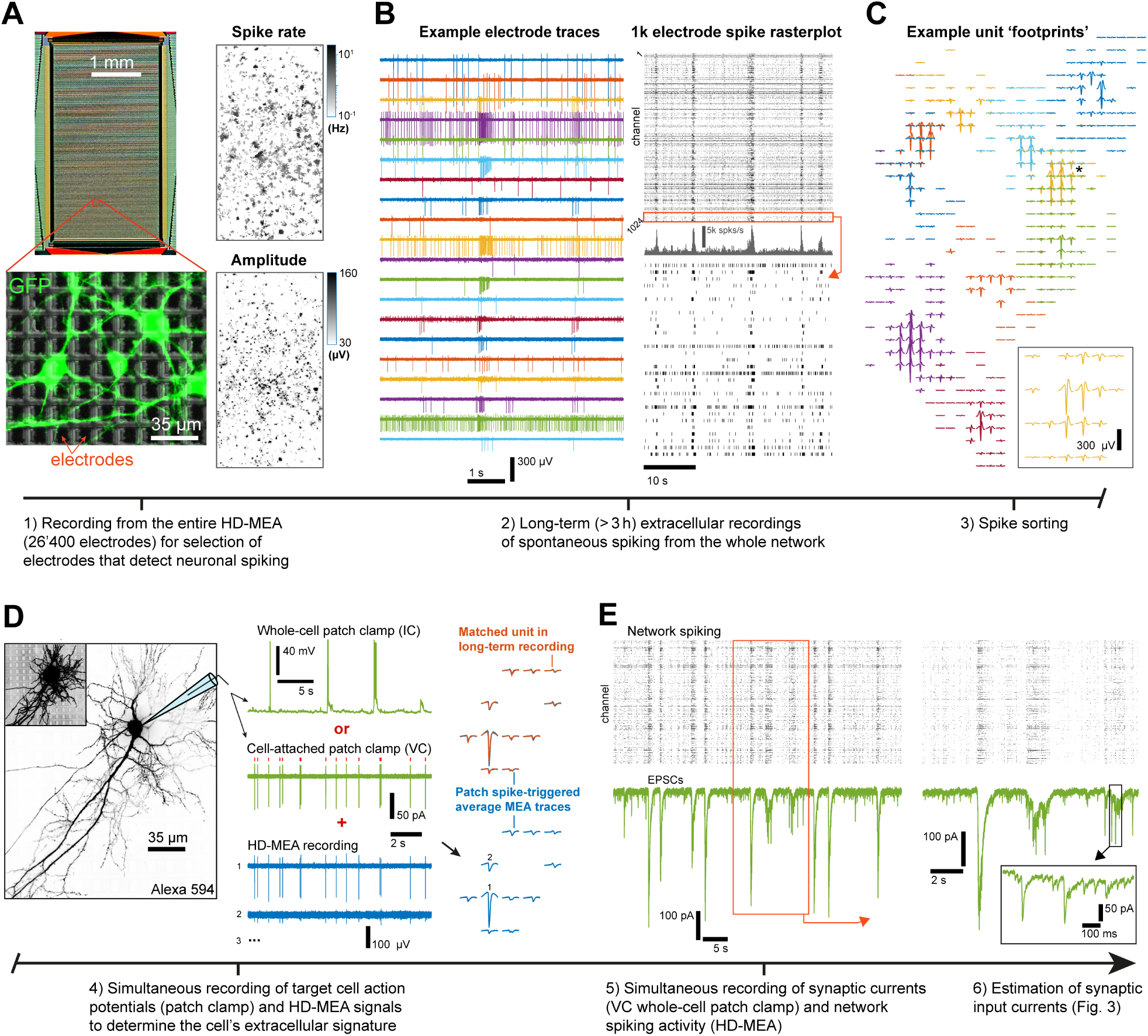
Experimental pipeline. **(A)** Left: Sensing area of a HD-MEA chip and magnified region with cultured primary neurons. Right: whole-array spike rate and amplitude maps. Electrodes displaying spiking activity were selected for subsequent long-term recording steps (grouped in 2-3 subsets of 1024 electrodes each). **(B)** Left: example electrode traces of a long-term recording. Right: spike raster plot of one entire electrode subset. The total recording time for each electrode was at least 3 h. **(C)** Example spike-triggered average extracellular signatures (or ‘footprints’) of individual units. A black asterisk marks the footprint magnified in the inset. Typically 100–200 units were identified per subset of 1024 electrodes. **(D)** Left: fluorescence images of a patched cell. Middle: example traces from a simultaneous patch-clamp (green) and HD-MEA (blue) recording. Right: spike-triggered averaging of the extracellular signals, based on spike times detected via the patch-clamp electrode, revealed the HD-MEA footprint of the patched cell (blue traces). The footprint of the patched cell was matched to a unit footprint from the preceding spike-sorted long-term recording (orange). **(E)** Electrode spike raster plot (top) and voltage-clamp trace (green; bottom) of a simultaneous HD-MEA and whole-cell patch-clamp recording. This paired recording was used to identify incoming monosynaptic connections (see Fig. 3). In all panels, HD-MEA electrode traces were band-pass filtered at 0.3–9.5 kHz.

In the final experimental step, we performed a second paired HD-MEA and patch-clamp recording to measure excitatory postsynaptic currents (EPSCs) in whole-cell voltage-clamp mode in addition to recording simultaneously extracellular network spiking (**Fig. 2E**). We used this second paired recording to estimate the average EPSC that was evoked in the patched cell by each of the extracellularly recorded neurons in the network – which enabled us to identify the neurons that were presynaptic to the patched cell. Note that a high-chloride internal patch-clamp solution was used and this shifted the chloride reversal potential. As a consequence of this shift, one voltage-clamp recordings at -70 mV holding potential was sufficient to simultaneously measure postsynaptic currents that were evoked by both glutamatergic (excitatory) and GABAergic (typically inhibitory) connections. Under such conditions, GABAergic cells also evoke net-inward currents (i.e., EPSCs) in the patched cell. We will later introduce a classification procedure for the connection type.

### Connectivity inference and EPSC estimation based on paired HD-MEA and patch-clamp recordings

We used the paired HD-MEA and whole-cell patch-clamp recordings from individual postsynaptic cells to identify all presynaptically connected neurons and to obtain an EPSC estimate for each incoming connection. Our method assumes that the patch-clamp current trace *I* at time point *t* is a linear superposition of the EPSCs of all neurons in the network:

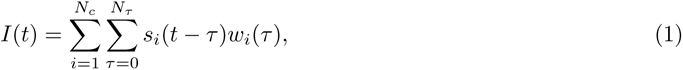

where *w_i_*is the EPSC waveform estimate (or, simply, EPSC) for the *i*^th^ neuron in the network, and *s_i_*(*t*) represents the corresponding (binary) spike train that indicates if neuron *i* spiked at time point *t*. *N_τ_* is the number of sample time points of the EPSC estimates. *N_c_* is the number of neurons in the network, which is the number of units resulting from spike sorting the HD-MEA data of the respective paired recording. Since we measured both the current trace (*I*) and the spike trains of the units in the network (*s_i_*), we can obtain the EPSC waveform estimates (*w_i_*) by linear regression. We adapted a previously published solution (Pillow et al. 2013) to this regression problem (see Methods for details), and upon its application to our paired recording data, a large number of synaptic EPSCs with a characteristic fast rising phase and a slow decay was identified (**Fig. 3**). In **Fig. 3A**, the monosynaptic connections and *w_i_* estimates are shown for one representative postsynaptic neuron. This example postsynaptic cell had 13 incoming connections with varying synaptic strengths and response onset latencies, while the > 100 remaining neurons/units in the network were putatively unconnected, as evident by their relatively flat EPSC estimate (i.e., these neurons did not show a significant evoked current).

**Figure 3.**
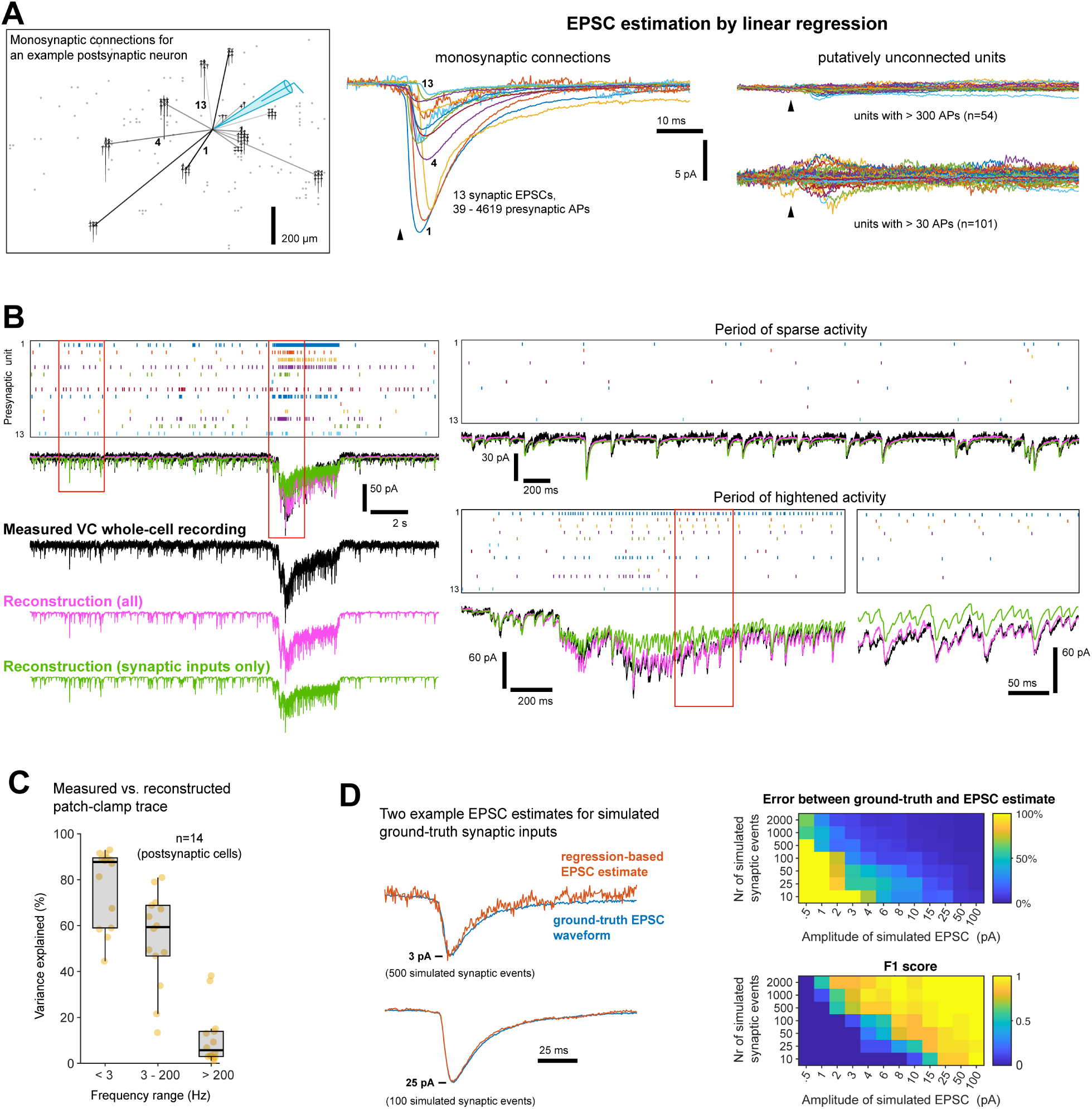
Connectivity inference and EPSC estimation based on paired HD-MEA and patch-clamp recordings. **(A)** Results from one example paired recording (e.g., Fig. 2E). We developed a linear regression procedure that estimates the EPSC that was evoked in the patched cell by each of the extracellularly recorded neurons. Middle: EPSCs of putative connections (amplitude > 10 × standard deviation of pre-spike baseline & amplitude > 1 pA). Black arrowhead indicates presynaptic spike time. Right: EPSC estimates of putatively unconnected units. Left: approximate spatial distribution of the presynaptic and postsynaptic cells. Locations of unconnected units are indicated by grey dots. **(B)** For the same paired recording as in (A), example recording period with a raster plot of presynaptic unit spiking (top), the VC patch-clamp recording (black), and two current-trace reconstructions using unit spike times and EPSC estimates of either all units (magenta) or of the putatively connected units only (green). **(C)** Variance of the measured patch-clamp recording explained by the respective current-trace reconstruction with all units (14 patched cells, with 142 identified connections, from 5 preparations). Different frequency contributions were compared (using fourth order Butterworth filters). Box plot indicates median and interquartile ranges, and whiskers the minimum/maximum values (excluding outliers). **(D)** Validation of the regression-approach by simulation of ground-truth synaptic inputs. Left: comparison of two example EPSC estimates with the corresponding ground-truth EPSC. Right: mean errors between ground-truth and EPSC estimate (top) and F1 score (bottom) for different simulation parameters (n = 14 simulations each; see Methods for details). See also Fig. S1 for evidence that the variation in the number of identified inputs is of biological origin.

To assess the EPSC estimation results, we performed reconstructions of the patch-clamp current trace based on the EPSC estimates and corresponding spike trains. The reconstruction was achieved by applying the right term of equation (1). Two different reconstructions were generated for each of the patched cells (**Fig. 3B**, same example cell as in Fig. 3A). For the first reconstruction (magenta), we used the *w_i_* and *s_i_* of all neurons/units in the network. For the second reconstruction (green), we only included the presynaptic cells forming a putative connection with the patched neuron. Note the often remarkable matching of measured (black) and reconstructed current traces. The two current-trace reconstructions were often very similar, indicating that the identified monosynaptic connections accounted for most of the observed currents. Some deviations were observed during periods of heightened activity, where especially slow currents played a role (e.g., see bottom-right traces in **Fig. 3B**).

Across a total of 14 patched putatively excitatory cells, 142 incoming connections were identified (mean = 10.1 *±* 5.2 s.d.; min = 3, max = 20 connections per cell). Excellent reconstruction performance was achieved across experiments. We separately compared slow baseline changes (< 3 Hz), fast synaptic activity (3 - 200 Hz) and putative high-frequency noise (> 200 Hz), yielding a median variance explained of approximately 60% in the 3 - 200 Hz range (**Fig. 3C**). Of note, some deviation were to be expected, e.g., due to synaptic transmission failures in the measured trace. Finally, we validated our regression approach by simulation of ground-truth synaptic inputs (**Fig. 3D**; see Methods for details). The simulation results suggested that our approach would only have failed to identify connections with extremely small-amplitude EPSCs and very low presynaptic spike rates.

In **Fig. 2D**, we described how the unit that corresponds to the patched cell can be identified in the long-term recording of network spiking that preceded the patch-clamp experiments by footprint matching. In a similar manner, we obtained parallel long-term spike trains of the presynaptic neurons that were identified by our regression method.

### Connection-type classification based on network-wide spike transmission or suppression

The usage of a high-chloride internal patch-clamp solution meant that both glutamatergic and GABAergic presynaptic cells evoked EPSCs in the patched cell, and, hence, a way to distinguish between the connection types was required. Here, we performed a connection-type classification by directly assessing if the respective presynaptic neuron had an inhibitory (i.e., suppressing) or excitatory (i.e., facilitating) effect on network-wide neuronal spiking. Using the > 3 h spike trains that were available for the presynaptic neurons, we computed the pairwise cross-correlograms (CCGs) and subsequently extracted the spike-transmission probability (STP) from each CCG (**Fig. 4A**; see Methods for details). The resulting STP measure is expected to be positive for excitatory connections and negative for inhibitory connections (Barthó et al. 2004). For cell-type classification, we computed a STP matrix based on all units that were recorded in parallel (**Fig. 4B**). A unit was then classified as putatively excitatory or inhibitory, if the mean (outgoing) STP value was positive or negative, respectively.

**Figure 4.**
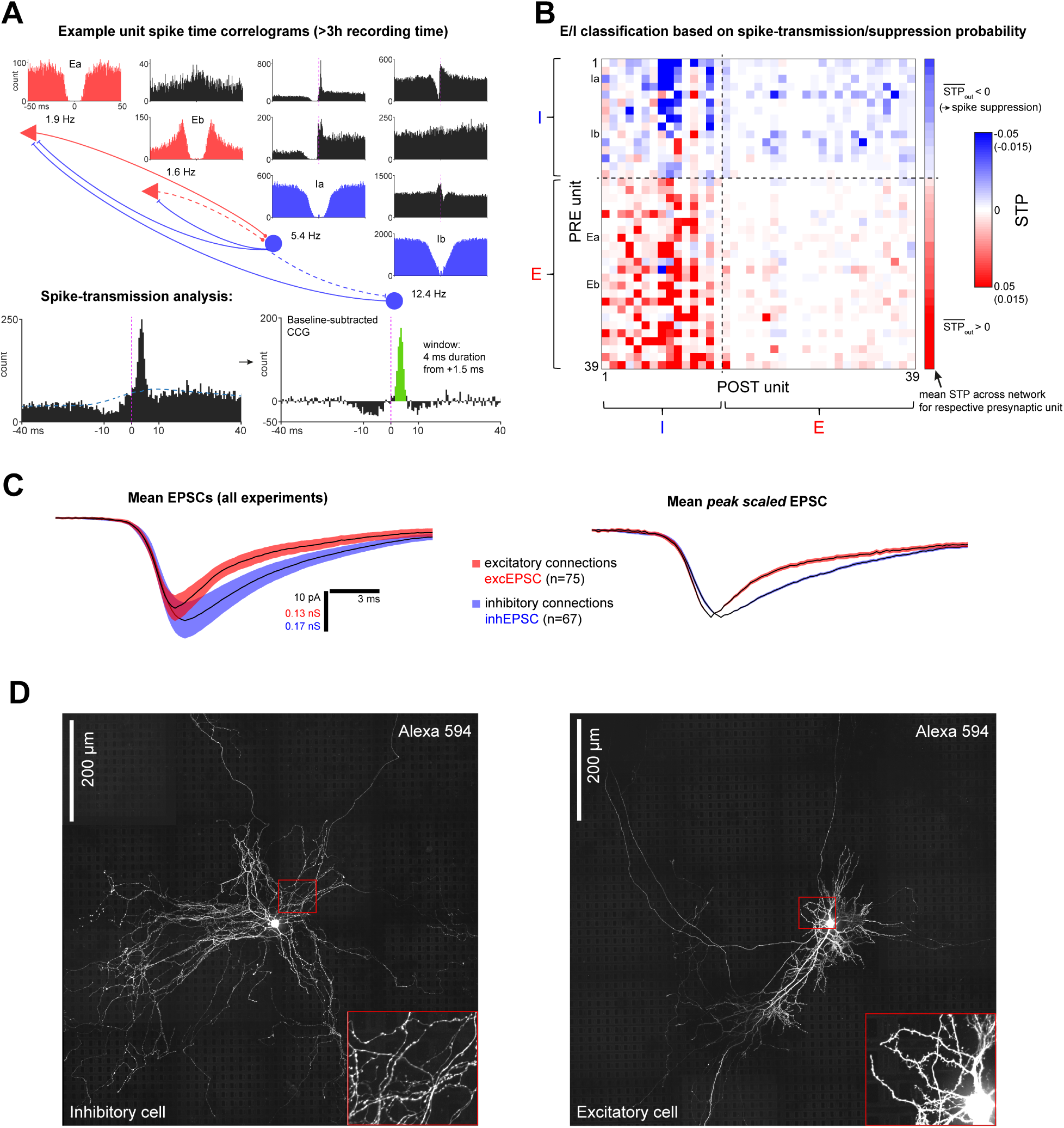
Connection-type classification based on network-wide spike transmission or suppression. **(A)** Top: auto- (red, excitatory cell; blue, inhibitory cell) and cross-correlograms (CCG; black) for four example units from a long-term HD-MEA recording (> 3 h). The schematic indicates putative connections. Bottom: illustration of spike-transmission probability (STP) extraction. The dashed blue line (left) indicates the CCG baseline. **(B)** Example matrix of STP values for all pairwise combinations of the units identified in a long-term HD-MEA recording. The rows labeled Ia/b and Ea/b correspond to the units shown in (A). The mean STP (across the sub-network) for each reference unit is shown on the right. Units are classified as being putatively excitatory or inhibitory based on the sign of their mean STP value. The color scale values in brackets apply to the column of mean STP values. Note the relatively large STP values associated with E-I compared to E-E unit pairs – in line with strong E-I connections typically found in the cerebral cortex (Campagnola et al. 2021). **(C)** Left: mean peak-aligned EPSCs, pooled from all experiments (14 patched cells) for connections that were classified to be excitatory (n = 75; decay *τ_fast_* = 2.5 ms, *τ_slow_* = 15.2 ms) or inhibitory (n = 67; decay *τ* = 8.6 ms). Right: same as on the left, but individual EPSC waveforms were first normalized with respect to their peak amplitude. Shadings denote the s.e.m. **(D)** Fluorescence images of a putative inhibitory and excitatory cell (Z-projections of stitched stack mosaics).

For 10 out of a total of 15 patched cells, the postsynaptic unit could be identified in the long-term HD-MEA recordings (see **Fig. 2D**); For the remaining 5, there was either no unit template available, or there was no matching unit found. Consistent with our attempt to target pyramidal cells, 9 out of the 10 cells with postsynaptic unit were classified as excitatory (mean STP > 0; mean spike rate = 1.2 Hz *±* 0.5 Hz s.d.) and only 1 cell was classified as inhibitory (mean STP < 0; spike rate = 9.6 Hz; cell subsequently excluded).

Following the cell-type classification, EPSC estimates could be attributed to an excitatory (‘excEPSC’) or inhibitory (‘inhEPSC’) connection. In line with known synaptic properties (Campagnola et al. 2021), the mean excEPSC across experiments exhibited faster kinetics compared to the mean inhEPSC (**Fig. 4C**). In this work, the term ‘EPSC’ refers to both excEPSCs and inhEPSCs. For a better comparison of excitatory and inhibitory synaptic activity, we also converted currents to conductances – separately for excEPSCs and inhEPSCs – based on the respective driving forces for glutamatergic and GABAergic ion channels. Note that this conversion expresses the effective conductance at the soma (the location of the patch-clamp electrode). In the following sections, we used the conductances for direct comparisons between excitatory and inhibitory inputs, and, otherwise, the exc/inhEPSC amplitudes.

Further supporting the validity of the classification approach, we also found striking differences in the extracellular footprint characteristics of the presynaptic neurons classified as excitatory and inhibitory, respectively (**Fig. S2**). Inhibitory cells exhibited a faster action potential propagation and larger footprint size, while excitatory cells had a more distant axonal projection. These footprint results were in agreement with fluorescence images of putative inhibitory and excitatory cells (**Fig. 4D**). The E/I classification concludes the main methodological part of this study.

### Spiking of individual neurons is dominantly controlled by a few strong incoming connections

After performing the analytical steps, detailed in the previous sections, we had identified multiple excitatory and inhibitory monosynaptic connections onto individual postsynaptic cells with corresponding estimates of the evoked postsynaptic currents. Moreover, long-term parallel spike trains of presynaptic and postsynaptic spiking activity were available (see **Fig. 1**). We exploited these data sets to investigate how synaptic activity relates to postsynaptic spiking. First, we focused on the role of individual monosynaptic connections, and, in particular, we examined the relationship between connection strength (i.e., EPSC amplitude) and the STP of the respective connection (**Fig. 5**).

**Figure 5.**
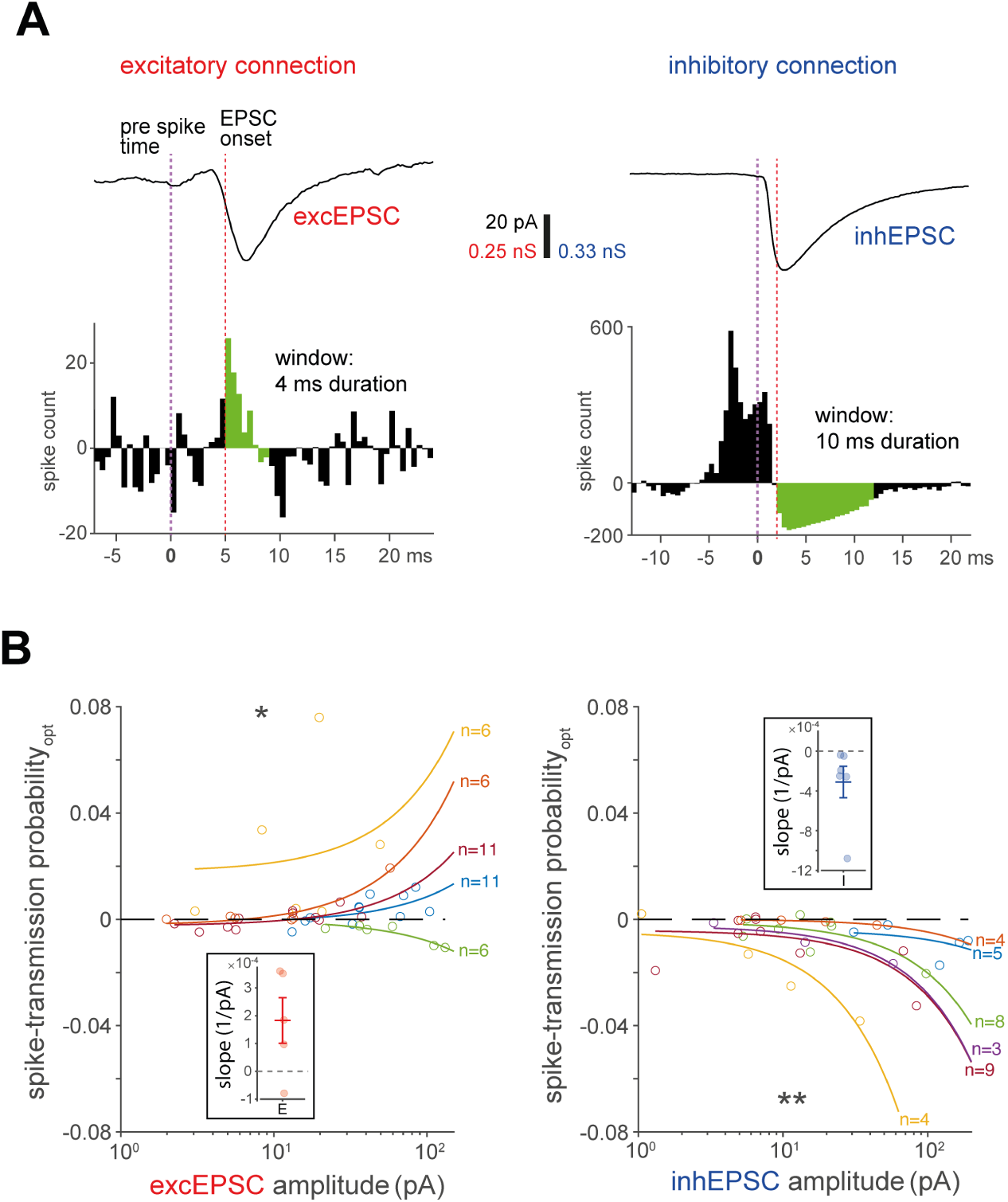
Spiking of individual neurons is dominantly controlled by a few strong incoming connections. **(A)** Optimization of STP estimates. Bottom: example baseline-subtracted CCGs, based on spike trains of a presynaptic and the postsynaptic unit. Top: EPSC of the corresponding connection. The EPSC onset latency relative to the presynaptic spike time was used to set the start time of the STP quantification window (green), thereby providing a more accurate STP estimate. This approach was particularly important for excitatory connections with variable excEPSC onset latencies; e.g., due to variable axonal length or synapse location within the dendritic tree. **(B)** Relationship between STP (optimized estimate) and EPSC amplitude for excitatory (left)/inhibitory (right) connections onto individual postsynaptic cells; lines are linear fits (here curves because of the logarithmic axis for amplitude; data points and linear fit belonging to the same postsynaptic cell are displayed in the same color). Insets show slopes from the linear fits with mean *±* s.d. Only cells with at least three E/I inputs were included ([E/I] 5/6 cells and 40/33 connections). Across all connections, the mean STP value for excitatory and inhibitory connections was 0.0025 *±* 0.0150 s.d. and -0.0090 *±* 0.0099 s.d., respectively. Data were comprehensively analyzed by linear mixed-effects modeling (with EPSC amplitude as a fixed effect and postsynaptic cell ID as a random effect). Significance was assessed by a likelihood ratio test. \**P* < 0.05, \*\**P* < 0.01

While STP has been suggested as a proxy for synaptic strength (English et al. 2017; Mizuseki and Buzsáki 2013), supporting experimental data is actually scarce. In fact, the influence of an individual connection on postsynaptic spiking is, besides connection strength, also strongly determined by the correlation of its activity with the activation of the other incoming connections. Varying input correlations and also variations in intrinsic neuronal properties could cause differences in the connection strength-STP relationship across different postsynaptic cells. To gain a clearer picture of the spike-facilitating or spike-suppressive effects of monosynaptic connections at postsynaptic-cell level, we calculated pairwise spike-time cross-correlations based on the spike trains of the presynaptic and postsynaptic neurons. As the time delay between presynaptic spike and postsynaptic exc/inhEPSC onset was known for each connection, we could align the STP quantification window to the response onset and, in this way, extract a further optimized STP estimate (**Fig. 5A**). For a comprehensive analysis that accounts for differences across postsynaptic cells, we used linear mixed-effects (LME) modeling (Yu et al. 2021), with the postsynaptic cell as a random effect (see Methods for details). This analysis indicated that, indeed, STP increased with increasing (absolute) ex-cEPSC amplitude of excitatory connections (0.011 per 100 pA *±* 0.012 s.e., *χ*^2^(1) = 6.4, *P* = 0.012; likelihood ratio test), and that STP decreased with increasing (absolute) inhEPSC amplitude of inhibitory connections (*−*0.031 per 100 pA *±* 0.010 s.e., *χ*^2^(1) = 6.8, *P* = 0.0089; likelihood ratio test). These results were consistent with individual linear regression fits, which were separately calculated based on the connection data of each postsynaptic neuron (**Fig. 5B**). The identified approximately linear relationship between STP and connection strength indicated that the degree of input correlation was typically similar across the connections received by individual postsynaptic cells during spontaneous recurrent activity. Two additional observations are worth pointing out. For one, there were considerable differences in regression-line slopes across the different postsynaptic cells, implying that connections of similar strength can have varying, cell-dependent effects on spiking. Furthermore, many of the (relatively weak) excitatory connections, remarkably, exhibited STP values scattered around zero, while inhibitory inputs had generally a more reliable (suppressive) effect on postsynaptic spiking. The results of this section indicate that the spiking of individual neurons was dominantly controlled by a few strong incoming connections, with a particularly important role for inhibitory inputs.

### Neuronal spiking is partially governed by rapid and brief changes in synaptic input activity

To elucidate the synaptic basis of postsynaptic spiking during spontaneous recurrent network activity, it is necessary to consider – in parallel – the combined excitatory and combined inhibitory conductances generated by the incoming connections. We reconstructed the synaptic activity experienced by the postsynaptic (patched) cells during the long-term extracellular recording period that preceded the patch-clamp experiments, as depicted in **Fig. 1** (see Methods for details). The observed alternations at the network level between periods of sparse spiking and periods of heightened, self-maintained network activity (see **Fig. 2B/E**) manifested as alternations between low and high conductance states (**Fig. 6A**). Synaptic conductances were often approximately balanced, with inhibitory conductance (*g_i_*) typically exceeding the excitatory conductance (*g_e_*), similar to estimations of the conductances generated by spontaneous network activity in vivo (Atallah and Scanziani 2009) and in vitro (Rudolph et al. 2007).

**Figure 6.**
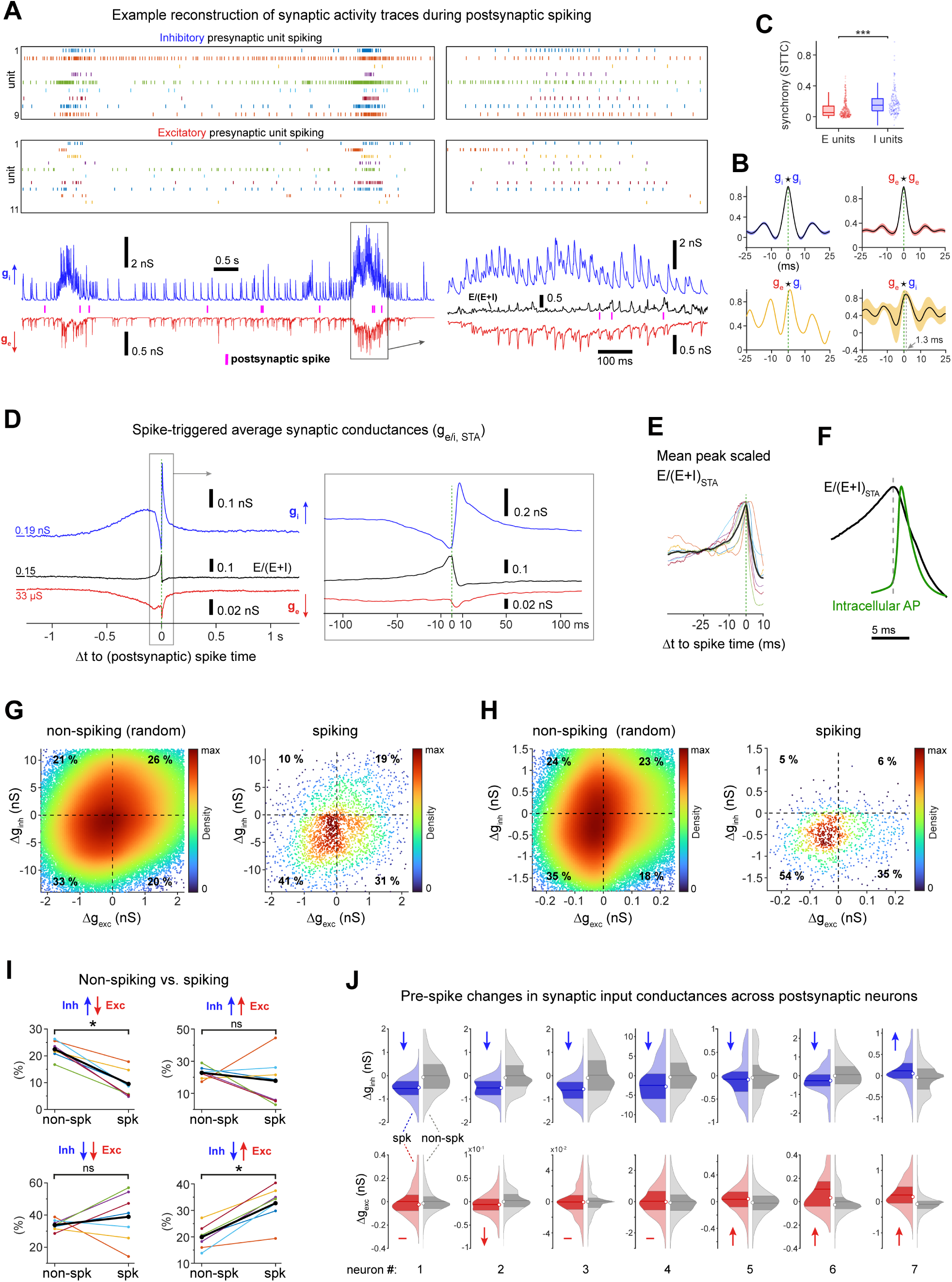
Neuronal spiking is partially governed by rapid and brief changes in synaptic input activity. **(A)** Top: raster plot of spike times for all presynaptic units of an example postsynaptic cell. Bottom: reconstructed inhibitory (*g_i_*; blue) and excitatory (*g_e_*; red) synaptic conductance traces. Spike times of the postsynaptic unit in magenta. *E/*(*E* + *I*) ratio (black trace; right). **(B)** Mean auto-(ACG) and cross-correlograms (CCG) of the reconstructed *g_i_*and *g_e_* of the neuron in (A) (25 individual mean-subtracted high-conductance events; shadings denote s.e.m.). **(C)** Pairwise unit synchrony (STTC with 10 ms binning [Cutts and Eglen 2014]; [E/I] 330/168 unit comparisons; *U* = 16288, *P* < 0.001, Mann-Whitney U test). Box plots indicate median and interquartile range, and whiskers the minimum/maximum values except for outliers. **(D)** Spike-triggered average of synaptic input conductances and *E/*(*E* + *I*) ratio of an example neuron (n = 15’813 postsynaptic spikes). See also Fig. S3 for variability across spikes and Fig. S4 for contributions of individual inputs. **(E)** Mean (black) and individual baseline-subtracted and peak scaled spike-triggered average *E/*(*E* + *I*) traces (n = 7 cells). **(F)** *E/*(*E* + *I*)*_ST A_* (black; mean from (E)) aligned to intracellular AP (green; mean from Fig. S5). See Fig. S5 for the temporal relationship between extracellularly measured spike times and intracellularly recorded action potential waveforms. **(G/H)** Density scatter plots of two example neurons, displaying the changes in input excitation and inhibition that occurred right before individual postsynaptic spikes (for each spike time t, we calculated the mean g value from t-1 ms to t+1 ms and subtracted the mean g value from t-20 ms to t-10 ms). As a reference (see plots labeled ‘non-spiking’), we also assessed the changes in input conductances that occurred during non-spiking periods (using random time points excluding periods *±* 20 ms from measured spikes; 10 x the number of spikes). **(I)** The percentage of postsynaptic spikes for the four possible pre-spike conductance-change combinations: increase in I and decrease in E (top-left; *Z* = 2.4, *P* = 0.018, Wilcoxon signed rank test); increase in I and E (top-right; *Z* = 0.68, *P* = 0.50); decrease in I and E (bottom-left; *Z* = -0.68, *P* = 0.50); decrease in I and increase in E (bottom-right; *Z* = -2.4, *P* = 0.018). Measured spikes (‘spk’) were compared to random time points from non-spiking (‘non-spk’) periods (n = 7 cells). **(J)** Violin plots of pre-spike changes in input inhibition (blue) and excitation (red) for individual postsynaptic neurons (n = 7). Distributions of non-spiking periods in grey. In each violin plot, the dark-colored area marks the interquartile range, the circle indicates the median, and the horizontal line marks the mean. All ‘spk’ vs. ‘non-spk’ comparisons significantly different except for excitation of neuron 1, 3, 4; otherwise P < 0.001 (Student’s t-test). Analyses from (G-J) focused on time periods with synaptic input activity (*g* > 2 × s.d. of *g* for both E and I). Across postsynaptic cells, 13’742 *±* 7’173 s.d. postsynaptic spikes were recorded, and 15% *±* 15% s.d. (min = 4%, max = 45%, median = 7%) occurred during high-activity periods. Some extreme values in (G-H) and (J) are not displayed. \**P* < 0.05, \*\**P* < 0.01, \*\*\**P* < 0.001.

The parallel reconstructions of the *g_e_* and *g_i_* traces also allowed us to calculate the E/I ratio during spontaneous network activity; here quantified as *E/*(*E* + *I*) (black trace, right, in **Fig. 6A**). Remarkably, this trace directly revealed brief spikes in the E/I ratio, during which postsynaptic action potentials preferentially occurred. We also assessed the temporal characteristics of the synaptic inputs in more detail (**Fig. 6B**). This analysis revealed, on average, a brief lag between excitation and inhibition and uncovered, particularly for inhibitory inputs, oscillatory dynamics, as observed in cortical and hippocampal networks in vivo (Atallah and Scanziani 2009; Buzsáki and Wang 2012; Okun and Lampl 2008; Salkoff et al. 2015). Moreover, quantifying the pairwise spike-train synchrony between all presynaptic cells showed a higher synchronization of inhibitory cells compared to excitatory cells (**Fig. 6C**).

To examine whether there were any major reoccurring synaptic events associated with postsynaptic spiking, we first generated the spike-triggered average of the inhibitory (*g_i,ST A_*) and excitatory input conductance (*g_e,ST A_*) for each postsynaptic cell. This analysis indicated that, on average, a sudden drop in inhibition often preceded postsynaptic spikes, followed by a rapid increase in inhibition, which was also reflected in the E/I ratio of the average conductances (**Fig. 6D**). For 7 out of 9 postsynaptic neurons, we found a fast increase in the input *E/*(*E* + *I*)*_ST A_* before postsynaptic spiking (**Fig. 6E**). We also uncovered the temporal relationship between the input *E/*(*E* + *I*)*_ST A_* and the intracellular action potential waveform (**Fig. 6F**). The aligned mean input-output data showed that the peak of the rapid E/I ratio increase coincided precisely with the action potential trigger time point, possibly indicative of a fine-tuned network organization.

For a more nuanced picture of which synaptic events are associated with postsynaptic spiking, we next quantified the changes in input excitation and inhibition that preceded individual postsynaptic spikes. In our analysis, we first focused on periods with high synaptic input activity. As previously discussed, cortical neurons in vivo typically receive and integrate barrages of input activation, similar to the high-activity events that we observed here (e.g., the event depicted in **Fig. 6A**, right). In **Fig. 6G/H**, individual pre-spike changes in input conductance are shown for two example postsynaptic neurons (plots labeled ‘spiking’, right). To assess how specific these conductance changes were to spiking periods, we also quantified the changes in input conductance that occurred during non-spiking periods as a reference (we used random time points from high-activity events excluding time points adjacent to measured spike times; we upscaled the number of measured spikes by 10×; the respective plots were labeled ‘non-spiking’). Spikes of both example neurons exhibited – compared to non-spiking regions – significantly more often a pre-spike decrease in inhibition, consistent with the mean conductance profiles. Precisely how an increase (top-right quadrants in **Fig. 6G/H**) or decrease (bottom-left quadrants) in both I and E conductance influenced the neuronal membrane potential is difficult to predict. However, if rapid changes in input conductance had a significant role in triggering spikes, one would expect that fewer spikes would exhibit a hyperpolarizing pre-spike increase in I and decrease in E (top-left quadrant) compared to the non-spiking period. Conversely, a decrease in I and an increase E (bottom-right quadrants) would likely result in a membrane potential depolarization so that more spikes should feature the corresponding pre-spike conductance changes compared to non-spiking periods. These relative shifts are precisely what can be observed in the plots of the two example neurons (**Fig. 6G/H**) and, in fact, across recordings (**Fig. 6I**). Finally, we compared the distributions of pre-spike changes in input inhibition and excitation of each postsynaptic neuron (**Fig. 6J**). Further indicating a pivotal role of inhibition in triggering spikes, 6 out of 7 neurons exhibited a clear decrease in the mean values (and medians) of pre-spike changes in inhibition compared to non-spiking periods. Interestingly, the 3 out of 7 neurons with an increase in excitation showed the smallest decrease in inhibition (or even an increase in inhibition in case of neuron #7). This latter observation suggests a matching of E and I inputs and cell-specific relative contributions of E and I conductance changes in triggering spikes.

Theoretically, neuronal spiking could be driven by a prolonged suprathreshold depolarization (Petersen and Berg 2016; Renart et al. 2007) or, in more favorable subthreshold regimes, by fast synaptic input fluctuations (Ahmadian and Miller 2021; Amit and Brunel 1997; Brunel 2000; Van Vreeswijk and Sompolinsky 1996). In this section, we demonstrated that the majority of investigated neurons featured – during high-activity periods – a significant number of spikes that were associated with rapid pre-spike changes in input conductances. These findings suggest that even simple neuronal cultures can self-organize to form circuits exhibiting sophisticated spiking dynamics.

### Network-state-dependent coordination of postsynaptic spiking by inhibitory inputs

Different network-activity levels presumably provide distinct means for the coordination of neuronal activity, with potential implications for the synaptic mechanisms of spiking. We, therefore, examined if the network state would influence the input conductances that were associated with postsynaptic spiking, and we focused on the dominant changes in inhibitory inputs. To this end, we first calculated the spike-triggered average conductances separately based on postsynaptic spike times that occurred either during high or low input conductance states (**Fig. 7**). Both low and high *g_i,ST A_* traces exhibited the characteristic bidirectional shape with a reduction in inhibition before the postsynaptic action potential, followed by a rapid increase in conductance (**Fig. 7A/B**). However, when comparing the temporal *g_i,ST A_* trace characteristics, it became apparent that the high *g_i,ST A_* conductance changes occurred considerably faster (**Fig. 7C**).

**Figure 7.**
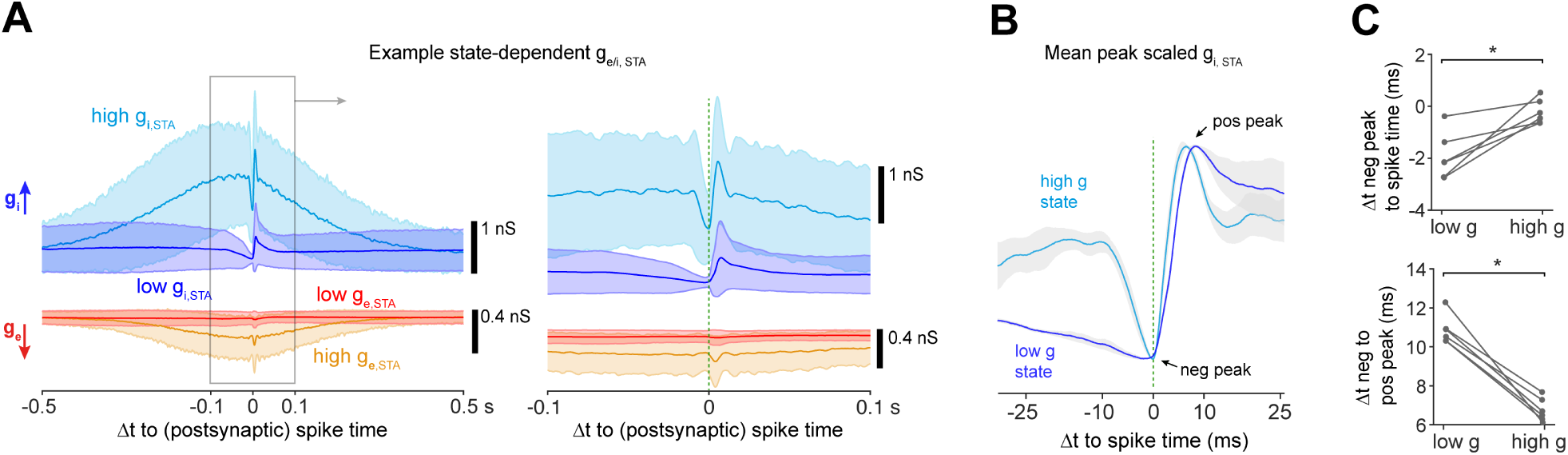
Network-state-dependent coordination of postsynaptic spiking by inhibitory inputs. **(A)** For an example postsynaptic cell, spike-triggered average synaptic conductances (*g_e/i,STA_*) based on postsynaptic spike times that occurred during periods of either high ([E/I] orange/light-blue) or low (red/blue) input conductance states (mean and s.d.). Right: Magnified region. Same cell as in Fig. 6D. **(B)** Mean *g_i,ST A_* across experiments (n = 6 postsynaptic cells; shadings denote s.e.m.) for high (light-blue) and low *g* (blue) states; scaled by their respective negative and positive peak values. Neurons with a pre-spike drop in inhibition were included in this analysis (neuron #1-6 from Fig. 6J). **(C)** Comparison of the time difference between the negative *g_i,ST A_* peak and postsynaptic spike time (*Z* = -2.2, *P* = 0.028, Wilcoxon signed rank test) and the time difference between the negative and positive *g_i,ST A_* peaks (*Z* = 2.2, *P* = 0.027) for low and high *g* states. Recording periods were labeled high *g* (conductance) state when *g_i_* > 3 × s.d. of *g_i_*, otherwise low *g* state. \**P* < 0.05.

How can these deviations in the mean conductance profiles be explained? To answer this question, we further quantified – separately for low and high g states – the changes in input inhibition that occurred right before and after individual postsynaptic spikes (**Fig. S6**). This single-spike analysis suggested that, during high g states, most spikes experienced a post-spike increase and pre-spike decrease in inhibition (see also **Fig. 6J**). On the other hand, low g states were characterized by sparse synaptic input (e.g., see reconstruction in **Fig. 6A**). Therefore, many of the spikes that occurred during low g states were associated with little change in input conductance (note medians of approximately zero in **Fig. S6A/C**). Nevertheless, a considerable fraction of spikes (often > 25%) from low g states were also associated with a post-spike increase and pre-spike drop in inhibition. It, therefore, appears that even the sparse inhibitory inputs of low g states could influence spike timing. Moreover, the post-spike increases in input inhibition during low g states suggest that there were strong regulatory inhibitory circuits in place. However, limited activity levels during low g states presumably introduced an increased jitter of these spike-associated changes in input inhibition.

In summary, the input inhibition of high-conductance states provides reliable and narrow windows-of-spiking opportunity. In addition, even during periods of sparse activity, there are rudimentary synaptic mechanisms in place to regulate spike timing.

### Organization of incoming monosynaptic connections at the level of individual postsynaptic cells

Having investigated the synaptic conductances that underlie the control of spike timing, we next examined the organization of the neural circuits that supported the observed spiking regime. Specifically, we characterized, from the perspective of individual postsynaptic cells, the distributions and relationships of key properties (synaptic strength, spike rate and EPSC onset delay) of the incoming monosynaptic connections (**Fig. 8**). A fundamental aspect of neural network organization is the fact that the distributions of network properties are often best characterized by a log-normal function, which has been linked to optimal information storage and processing principles (Barbour et al. 2007; Buzsáki and Mizuseki 2014). First, we tested if the methods developed in the previous sections could reveal similar organizational principles in our neuronal culture model. We pooled the connection data from all recordings; and indeed, the distributions of all properties, with the exception of inhibitory onset latency, were well described by a log-normal distribution (left panels in **Fig. 8B-D**). Moreover, all properties showed significant differences between excitatory and inhibitory connections.

**Figure 8.**
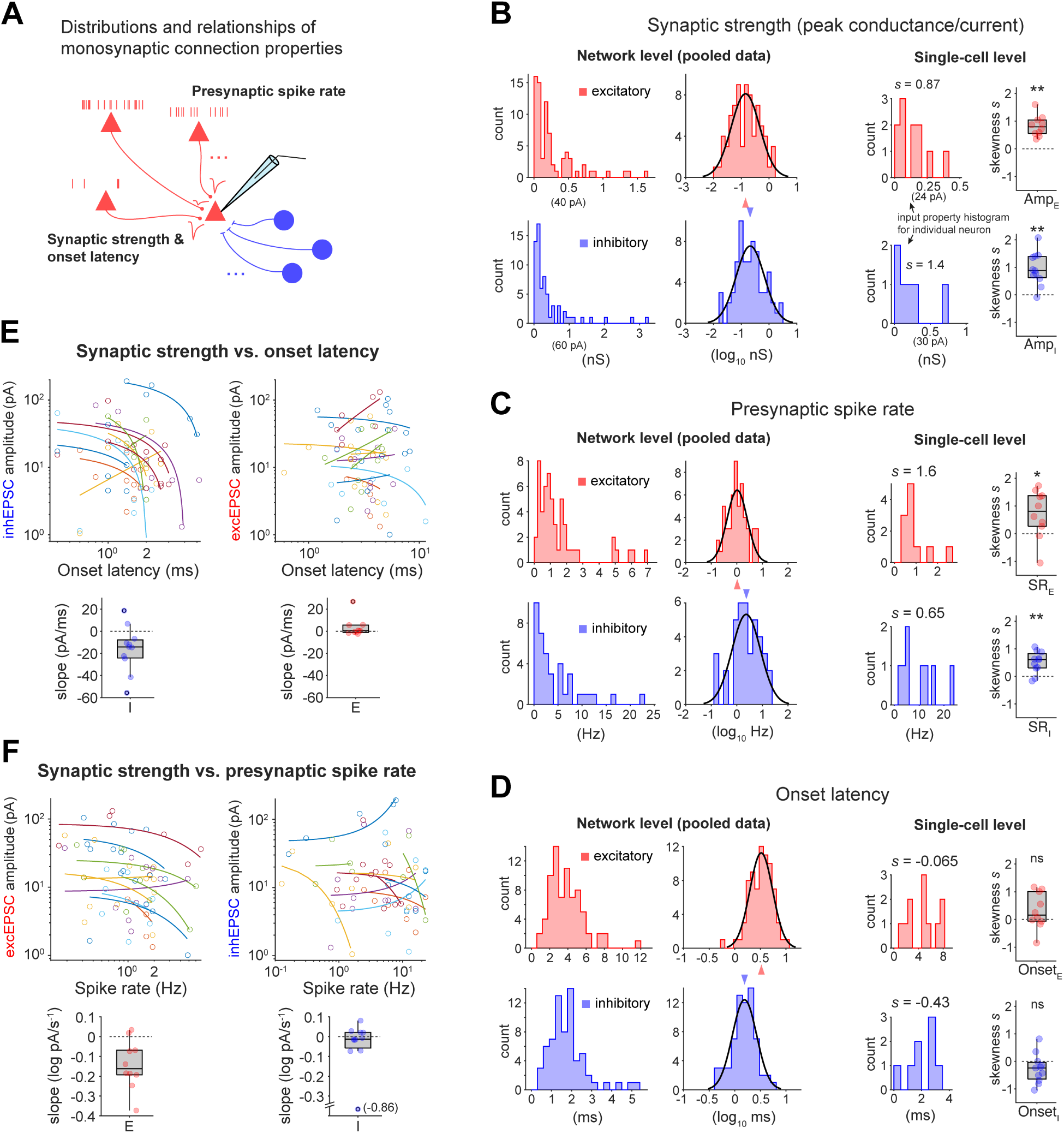
Organization of incoming monosynaptic connections at the level of individual postsynaptic cells. **(A)** Presynaptic spike rate, amplitude of the EPSC estimate (also converted to conductance) and EPSC onset latency were analyzed. **(B-D)** For each property, data were examined at the network level (connections from all 14 paired recordings pooled) and from the perspective of the individual postsynaptic cells. For the network-level analysis (left), a histogram with and without previous log-transformation are shown (black curve: Gaussian fit; arrow head marks the peak). For all properties, including amplitude (*U* = 1660, *P* < 0.001, n[E/I] = 75/67), spike rate (*U* = 601, *P* < 0.001, n[E/I] = 45/33) and onset latency (*U* = 739.5, *P* < 0.001 n[E/I] = 75/67), there was a significant difference between excitatory and inhibitory connections (Mann-Whitney U test). Moreover, all distributions, except for that of inhibitory onset latency (D), were approximately log-normal (Shapiro-Wilk test). For the single-cell-level analysis (right), an example single-cell histogram is shown in addition to a box plot of the skewness *s*. The skewness of amplitude ([E/I]; *Z* = 2.8/2.8, *P* = 0.0051/0.0044, n = 10/11 postsynaptic cells) and spike rate (*Z* = 2.1/2.7, *P* = 0.037/0.0076), but not onset latency (*Z* = 1.4/-1.5, *P* = 0.17/0.13), was significantly different from zero (Wilcoxon signed rank test). **(E)** Relationship between EPSC amplitude and onset delay at single-cell level. Top: scatter plot, with one linear regression fit for each postsynaptic cell. Bottom: slope values of linear fits. **(F)** Relationship between EPSC amplitude and presynaptic spike rate at the single-cell level. Amplitude values were log-transformed, otherwise as in (E). Only cells with at least 3 E/I inputs were included (n[E/I] = 10/11 cells). Data in (E/F) were comprehensively analyzed by linear mixed modeling. Box plots indicate median and interquartile ranges and whiskers the minimum/maximum values except for outliers. \**P* < 0.05, \*\**P* < 0.01

Consistent with typical in vivo findings, inhibitory cells displayed higher spike rates and conductances and lower onset latencies compared to excitatory cells, indicative of a relatively strong and fast action of inhibition. The fast inhibition (i.e., small onset latencies) was likely the result of both a relatively local inhibitory innervation and a fast axonal action potential propagation (**Fig. S2**).

How are the properties of incoming connections organized from the perspective of individual postsynaptic cells? Are distributions skewed, as observed at the network level, or more homogeneous? Even though the nature of these property distributions would provide important insights into neural functioning, experimental single-cell data and corresponding characterizations are scarce. Next, we therefore used our data sets to calculate the skewness *s*, for each postsynaptic cell and each incoming connection property (right panels in **Fig. 8B-D**). Synaptic strength and spike rate were typically associated with a positively skewed distribution at the single (postsynaptic)-cell level. These results suggested that individual neurons were particularly strongly influenced by a few key inputs, while the majority of incoming connections played – individually – a relatively small role.

Finally, we examined – at the level of individual postsynaptic cells – the relationships between connection strength (i.e., absolute EPSC amplitude) and either the EPSC onset latency or the presynaptic spike rate. Uncovering these relationships may provide indications of the regulatory processes of synaptic strength that were implemented by the networks. We fitted individual linear regression lines for each postsynaptic cell (**Fig. 8E/F**) and performed a comprehensive analysis across cells using linear mixed-effects (LME) modeling. We found that for most postsynaptic cells, the strength of incoming inhibitory connections decreased with increasing onset latency, and the relationship was well characterized by linear regression fits (**Fig. 8E**; according to LME model: -14.2 pA *±* 2.8 s.e. per ms, *χ*^2^(1) = 13.1, *P* < 0.001; likelihood ratio test). This result provided further evidence that individual postsynaptic cells were dominated by strong local inhibition. Furthermore, the strength of excitatory connections decreased with increasing presynaptic spike rate, and the relationship was well described by an exponential decay (**Fig. 8F**; amplitude values were linearized by log-transformation; according to LME model: -8.4 % *±* 3.2 s.e. decrease in amplitude per Hz, *χ*^2^(1) = 4.8, *P* = 0.029). This finding indicates a homeostatic synaptic plasticity mechanism so as to achieve a downregulation of synaptic strength for connections that were particularly active.

### A few key inhibitory hub cells with high spike rates, strong synapses and fast action potential propagation dominate the network

The results of the previous sections showed that individual neurons were particularly strongly influenced by a few connections with strong synapses (**Fig. 8**), which translated to strong effects on postsynaptic spiking (**Fig. 5**). Did the presynaptic neurons that provided these important inputs exhibit characteristic properties or were these inputs of random neuronal origin? To answer this question, we characterized the organizational principles concerning outgoing connections (**Fig. 9**). We focused on a network, in which multiple paired HD-MEA and patch-clamp recordings were sequentially obtained from different postsynaptic cells (**Fig. 9A**). Following the identification of incoming connections for each paired recording using our regression approach, we found that the same presynaptic cell formed often connections with multiple postsynaptic (patched) cells (see matching footprints in top inset in **Fig. 9A** and **Fig. S7**). There was a variety of different outdegrees, with relatively few highly connected presynaptic cells (**Fig. 9B**). Individual presynaptic cells evoked EP-SCs with drastically varying amplitudes in different postsynaptic cells (bottom inset in **Fig. 9A**), and the distribution of outgoing connection amplitudes was typically positively skewed (**Fig. 9C**). Next, we asked whether there was a relationship between the degree of outgoing connectivity and other neuronal properties that determine the cell’s influence on the network. Indeed, for inhibitory – but not excitatory – cells, the spike rate of a neuron correlated with its outdegree (**Fig. 9D**). Furthermore, the sum of outgoing inhEPSC amplitudes of individual inhibitory cells increased rapidly with the outdegree, while for excitatory cells this increase was slower (left in **Fig. 9E**). To examine if the degree of outgoing connectivity influences the mean connection strength, we first calculated the mean postsynaptic EPSC waveform (*EPSC_out_*) for each presy-naptic cell. Subsequently calculating the mean waveform of all *EPSC_out_* from cells with outdegree 1-3 and 4-7, respectively, indicated that highly connected inhibitory – but not excitatory – cells formed particularly strong synapses with their postsynaptic targets when compared to cells with few outgoing connections (right in **Fig. 9E**). Finally, the action-potential propagation velocity of inhibitory cells (see Methods for details) also increased with the outdegree (**Fig. 9F**). In summary, a few key inhibitory cells were in a unique position to coordinate network activity by exerting fast and strong effects through an extensive network of outgoing connections.

**Figure 9.**
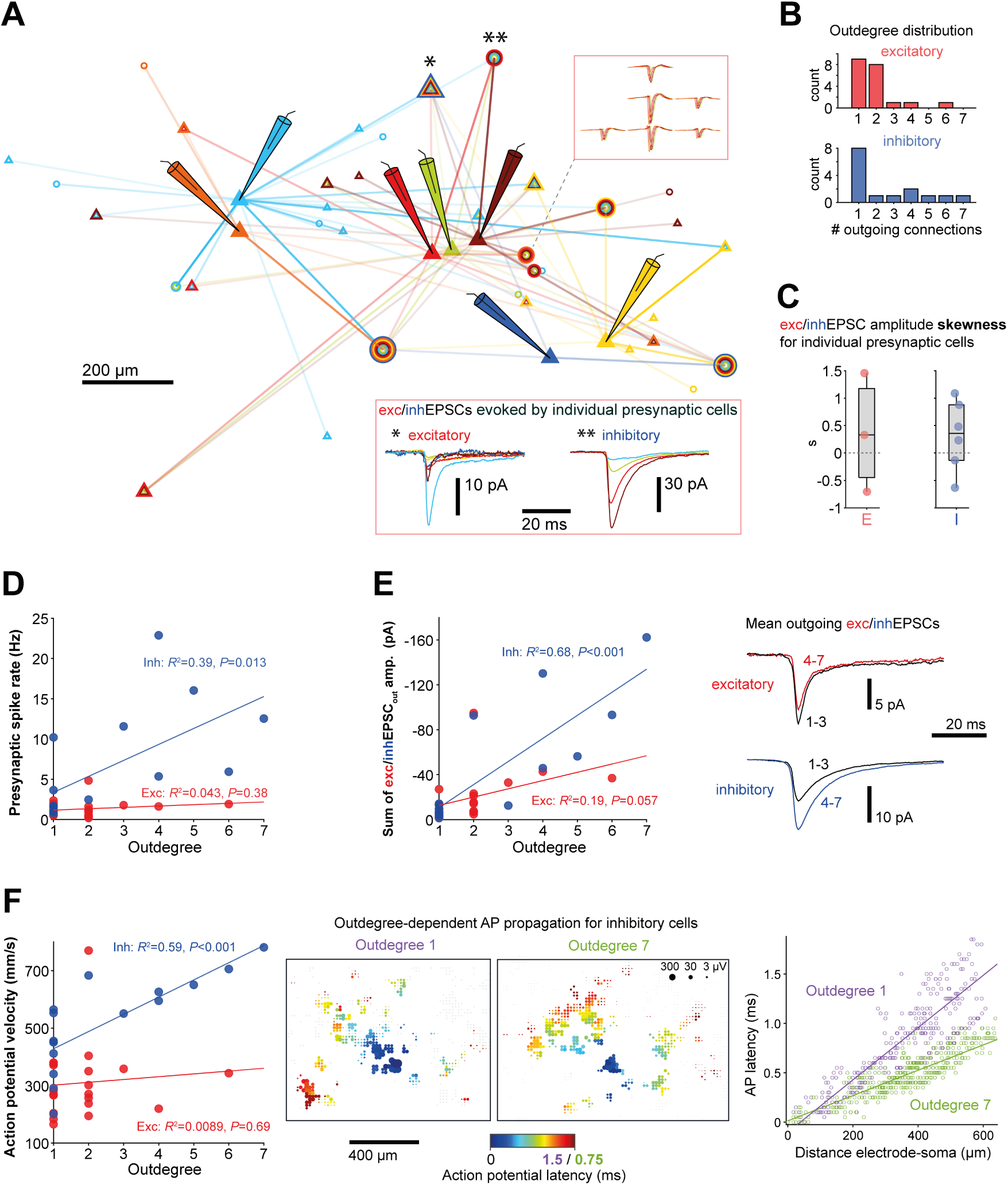
A few key inhibitory hub cells with high spike rates, strong synapses and fast action potential propagation dominate the network. **(A)** Multiple neurons in the same network were sequentially patched (locations approximated by electrode schematic). Paired patch-clamp and HD-MEA recordings were performed for each cell, followed by identification of their incoming connections using our regression approach. Open triangles and circles mark the positions of excitatory and inhibitory cells, respectively. Overlaid circles/triangles represent neurons that were presynaptic to multiple patched cells. Top inset: virtually identical example extracellular footprints from four separate paired recordings (traces slightly shifted in time for better visibility). Bottom inset: Example EPSCs evoked in different postsynaptic cells by the same excitatory/inhibitory presynaptic cell (marked in the network schematic by single/double asterisk). **(B)** Outdegree distribution for the network shown in (A). **(C)** Skewness *s* of the amplitude distribution for the outgoing connections of individual neurons (minimum of 3 connections). **(D)** Relationship between the spike rate of a neuron and its outdegree. Significance was assessed after applying the Holm correction for multiple comparisons. **(E)** Left: Relationship between the sum of EPSC amplitudes of all outgoing connections of a neuron and its outdegree. Right: for each presynaptic cell, the mean postsynaptic EPSC waveform 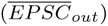 was computed; the mean of all 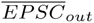 from cells with outdegree 1-3 and 4-7 are shown. **(F)** Left: Relationship between action potential velocity of a presynaptic neuron and its outdegree. Middle: Example HD-MEA action potential (AP) latency plot from a neuron with outdegree 1 and 7. Right: Plots of AP latency vs. distance between electrode and soma for the two example units.

## Discussion

We examined the synaptic mechanisms that determine postsynaptic spike timing during spontaneous recurrent network activity – linking functional and organizational circuit characteristics. These investigations were made possible through several methodological innovations including a comprehensive synaptic input mapping in conjunction with long-term extracellular whole-network recordings, which allowed for reconstructing synaptic activity during a period of postsynaptic spiking. For the input-mapping approach, based on simultaneous HD-MEA and patch-clamp recordings, we developed and validated a regression procedure that inferred a large proportion of the (current-evoking) monosynaptic connections onto individual postsynaptic cells. Compared to even the most advanced multi-channel patch-clamp platforms (Campagnola et al. 2021; Peng et al. 2019), our approach exceeded the number of testable incoming connections by an order of magnitude. To achieve good network coverage, neurons needed to be located adjacent to an electrode, which puts constraints on the possible network size. While the present study already made use of modern CMOS HD-MEA technology, new generations of HD-MEAs will feature further increased electrode densities, larger recording areas, and larger sets of electrodes that can be recorded from simultaneously (Dragas et al. 2017; Yuan et al. 2020). Thus, utilizing next-generation HD-MEAs with our approach will immediately improve network coverage and allow for monitoring even larger neuronal networks. In this study, we chose to examine in vitro circuits, formed in neuronal cell cultures, as modern in vitro HD-MEAs can provide near-complete network coverage. In addition, HD-MEA devices have been successfully combined with patch clamping in vivo (Marques-Smith et al. 2018), and the strategies and software tools introduced in our study will be equally applicable in an in vivo context.

During sensory stimulation and spontaneous network activity, neurons typically experience a proportional or ‘balanced’ change in excitatory and inhibitory inputs (Haider and McCormick 2009; Isaacson and Scanziani 2011; Liu 2004; Shu et al. 2003). Theory suggests that multiple dynamical regimes could yield this balance (Brunel 2000; Renart et al. 2007), and understanding the nature of the implemented regime is crucial for gaining a better understanding of cortical function in general. Theoretical and computational modelling studies further indicate that, during the balanced state, rapid membrane potential fluctuations could be the primary spike trigger (Ahmadian and Miller 2021; Amit and Brunel 1997; Brunel 2000; Van Vreeswijk and Sompolinsky 1996). These studies typically assume randomly connected networks with asynchronous irregular firing. It remains to be seen to what extent these findings hold true in biological networks with structured connectivity and – as often seen during spontaneous cortical activity – more synchronous spiking (Ahmadian and Miller 2021). In addition, other dynamical regimes have been described, including spiking driven by the mean input rather than rapid input fluctuations (Petersen and Berg 2016; Renart et al. 2007). Here, we directly tested in biological neural networks if postsynaptic spiking was associated with rapid changes in synaptic inputs. First, we showed that, for individual postsynaptic cells, the strength of excitatory and inhibitory connections correlated with the degree of postsynaptic-spike transmission and suppression, respectively. However, many of the spike-time cross-correlograms associated with excitatory connections did not show clear peaks of excess spiking, which may indicate that these inputs provided more of a ‘basal excitatory tone’ and, hence, did not consistently influence postsynaptic spike timing. On the other hand, the spike-suppressive effect of inhibitory inputs was found to be generally reliable. In line with these results, the reconstruction of synaptic input conductances revealed that postsynaptic spiking often coincided with the peak of E/I ratio fluctuations. These changes in E/I ratio were – depending on the postsynaptic cell – primarily due to either an increase in excitation or decrease in inhibition, while the decrease in inhibition outweighed the increase in excitation in most cells. The observed ability to respond to different combinations of conductance changes is consistent with reports of dynamic-clamp experiments (Piwkowska et al. 2008). We also showed that a rapid increase in inhibition typically restricts the spiking window and that the temporal characteristics of inhibition around the postsynaptic spike time is sharpened during periods of heightened network activity. The latter observation could be the result of an increased synchronization of inhibitory neurons (Buzsáki and Wang 2012; Destexhe et al. 2001; Hasenstaub et al. 2005). In summary, we found that the studied biological networks were equipped with several mechanisms that allowed them to operate, at least partially, in a precise dynamical regime governed by rapid input changes. Interestingly, cortical neurons in vivo typically receive – in response to diverse sensory stimuli – stereotypical excitation-inhibition sequences and, in this scenario, the excitatory inputs presumably determine the initial timing of the depolarization (Monier et al. 2003; Okun and Lampl 2008; Wehr and Zador 2003). To consolidate these findings with the observed role for inhibition in controlling spike timing, it is conceivable that there are two main modes of neuronal spiking: external inputs trigger a depolarization due to an increase in excitation, while spontaneous spiking (e.g., associated with slow-wave sleep or neuronal cell-culture activity) is predominantly driven by local recurrent activity and, in particular, by a reduction in inhibition. It is an intriguing question to ask what the implications of these different spiking modes would be (e.g., for spike-timing dependent plasticity). Finally, we investigated the circuit architecture that supported the observed spiking regime. Highly connected hub neurons are believed to be important for the coordination of network activity (Bonifazi et al. 2009; Cossart 2014; Gal et al. 2021), but electrophysiological characterizations of their connections are scarce. Here, we also identified a minority of cells with high connection outdegrees. Moreover, we found that inhibitory cells that were highly connected were also fast-spiking and featured relatively strong synapses and fast action potential propagation velocities. In addition, when we examined the incoming connections onto individual postsynaptic cells, we found that connection strength and presynaptic spike rate followed approximately a log-normal distribution, implying that a few incoming connections were particularly dominant. Combining these findings on the organization of incoming and outgoing connections with our functional data, a picture emerges in which network-wide neuronal spiking is effectively coordinated by a few key inhibitory hub neurons with windows of spiking opportunity provided by a brief reduction in their postsynaptic effects.

The findings presented in this work provide a detailed characterization of a dynamical regime that is in line with theoretical predictions for neural networks in vivo. Our results were obtained in a cortical cell culture model and we, therefore, propose that a self-organization towards a dynamical regime governed by rapid input changes is an inherent property of cortical networks.

## Materials and Methods

### Primary neuron culture preparation

The experimental protocols involving animal tissue harvesting were approved by the veterinary office of the Canton Basel-Stadt according to Swiss federal laws on animal welfare and were carried out in accordance with the approved guidelines. Before cell plating, the HD-MEA chips were sterilized for 45 min in 70% ethanol and washed 3 × with sterile deionized (DI) water. Next, the electrode array was treated with 20 *µ*L of 0.05% (v/v) poly(ethyleneimine) (Sigma-Aldrich) in borate buffer (Thermo Fisher Scientific) at 8.5 pH, for 40 min at room temperature, and then washed 3 × with DI water. Subsequently, we added 8 *µ*L of 0.02 mg mL*^−^*^1^ laminin (Sigma-Aldrich) in Neurobasal medium (Gibco) and incubated the chips for 30 min at 37 °C. Cortices of E-18 Wistar rat embryos were harvested in ice-cold HBSS (Gibco) and then dissociated in trypsin with 0.25% EDTA (Gibco). We next seeded 15’000 to 20’000 cells on top of the electrode array. Subsequently, the chips were incubated at 37 °C for 30 min before adding 2 mL of plating medium. The plating medium stock solution consisted of 450 mL Neurobasal (Gibco), 50 mL horse serum (HyClone, 1.25 mL Glutamax (Invitrogen), and 10 mL B-27 (Invitrogen). Every 3-4 days, 50% of the culture medium was replaced by growth medium, with the stock solution consisting of 450 mL D-MEM (Invitrogen), 50 mL horse serum (HyClone), 1.25 mL Glutamax (Invitrogen), and 5 mL sodium pyruvate (Invitrogen). The HD-MEA chips were kept inside an incubator at 37 °C and 5% CO_2_. All the experiments were conducted between days in vitro (DIV) 15-18, when cellular growth and network connectivity had stabilized.

### High-density microelectrode array (HD-MEA) system

A complementary-metal-oxide-semiconductor (CMOS)-based HD-MEA featuring 26’400 electrodes (pitch of 17.5 um) within an overall sensing area of 3.85 × 2.10 mm^2^ was used (Müller et al. 2015). An arbitrary subset of these electrodes could be connected to 1024 channels for simultaneous readout at 20 kHz sampling frequency. The HD-MEA system was developed in-house, but can also be purchased as the MaxOne model (MaxWell Biosystems). The electrodes were coated with electrodeposited platinum black to decrease electrode impedance and improve the signal-to-noise characteristics.

### Electrode selection and long-term extracellular recording of network spiking

To select the HD-MEA electrodes for long-term recordings, all 26’400 electrodes were initially briefly scanned for activity (1 min per electrode). Electrodes that recorded spiking activity (minimum electrode spike rate: 0.05 Hz) were then identified and ranked according to their mean spike amplitude. Next, 2 or 3 × 1024 of the electrodes with the largest signals were selected, which typically covered most of the active network. The selected electrodes were then divided into sets of up to 1024 electrodes along the longitudinal axis of the chip for simultaneous read-out (adjacent electrode sets had a small overlap of 3 electrodes width). To obtain long-term recordings of network spiking, we sequentially recorded from the electrode sets for 15 min each and repeated this every 1 h for a total recording time of at least 3 h for each electrode set.

### Patch-clamp electrophysiology

For simultaneous patch-clamp and HD-MEA recordings, cultures were transferred immediately after the long-term network recording period from the incubator to a patch-clamp setup with integrated HD-MEA recording unit. From the the electrode sets that were used for long-term recordings, we then typically selected the electrode set covering the largest chip area with an even cell distribution and targeted pyramidal-shaped neurons towards the center of the electrode set for patching. The patch-clamp setup comprised a MultiClamp 700B amplifier (Axon Instruments) and an Axon Digidata 1440A (Axon Instruments). Data were low-pass filtered at 5 kHz and sampled at 20 kHz, with data acquisition controlled by the sofware WinWCP. Synchronization pulses were generated via the Digidata unit and fed into the HD-MEA system for post-recording data alignment. Cells were perfused with BrainPhys^TM^ Neuronal Medium (Stem Cell Technologies) heated to approximately 32-34°C. Cell-attached and whole-cell patch-clamp recordings in voltage-clamp mode were obtained with standard borosilicate glass micropipettes (4–5 MΩ) containing the following internal solution (in mM): 85 caesium-gluconate, 60 CsCl, 10 Hepes, 4 Na_2_ATP, 0.3 GTP, 2 MgCl_2_, 0.1 EGTA, (pH 7.2- 7.3; 280–290 mOsmol/l). Alexa Fluor 594 (20 *µ*M) (Sigma-Aldrich) was added for cell morphology assessment. The holding potential was set to -70 mV (without liquid-junction potential [LJP] correction). Only cells with a series resistance smaller than 25 MΩ were included in our study. We chose an internal solution with a relatively high chloride concentration, which causes a polarity reversal of GABA-A receptor-associated currents due to a more positive reversal potential (similar to early developmental periods). This approach allowed us to simultaneously record postsynaptic currents evoked by both GABAergic and glutamatergic synapses. We calculated the chloride reversal potential to be approximately -20 mV using the Nernst equation. Therefore, we used driving forces of 60 mV (inhibition) and 80 mV (excitation) to convert the synaptic currents to conductances according to Ohm’s law (with a -10 mV LJP correction). The fact that both GABAergic and glutamatergic presynaptic neurons evoked EPSCs in the patched cells meant that a way to distinguish between the connection types was required. Typical excitatory-inhibitory classification strategies for 3D tissue, based on unit extracellular signatures and spiking behavior (Barthó et al. 2004; Csicsvari et al. 1998; Senzai and Buzsáki 2017), appear to be not always sufficient for 2D preparations (Weir et al. 2015). We, therefore, performed a connection-type classification by assessing if a given neuron’s activity is typically associated with spike transmission or suppression.

For paired IC patch-clamp and HD-MEA recordings to examine the relationship between extracellular and intracellular action potentials and for dye loading to image neurites at high-resolution, the following internal solution was used (in mM): 110 potassium-gluconate, 10 KCl, 10 Hepes, 4 MgATP, 0.3 GTP, 10 phosphocreatine, (pH 7.2-7.3; 280–290 mOsmol/l). On the day of the experiment, Alexa Fluor 594 (20 *µ*M and 50 *µ*M for paired recordings and imaging experiments, respectively) was added.

To generate the spike-triggered average HD-MEA footprint of the patched cell, we used spontaneous spiking recorded in cell-attached mode or during a brief IC whole-cell recording. Additional spikes were sometimes triggered via current injection to increase the total number of spiking events.

### Confocal fluorescence microscopy

A Nikon NiE upright confocal microscope featuring a Yokogawa W1 spinning disk, an ORCA-Flash4.0 V2 Digital CMOS camera (Hamatsu Photonics), and a 60x/1.00 NA water-objective (Nikon) was used for fluorescence imaging. To generate large field-of-view fluorescence images of neurite projections at high-resolution, individual neurons were loaded with Alexa Fluor 594 via the patch-pipette for at least 30 min. Multiple imaging tiles covering most of the cell morphology were defined. For each tile, a z-stack of images was acquired (0.4 µm z-step; 0.1125 µm x-y resolution). Using Huygens Professional (version=21.10; Scientific Volume Imaging), images were first deconvolved (CMLE algorithm) and then stitched together (10% overlap, circular vignetting correction model). A 561 *nm* excitation laser was used in combination with a 609*/*54 *nm* emission filter. To image the distribution of neurons on the HD-MEA chips, cultures were transduced with floxed EGFP (AAV9/2-hSyn1-chI-loxP-EGFP-loxP-SV40p(A); MOI = 5x10^5^ vg) and Cre (AAV9-hSyn-Cre-WPRE-hGH; MOI = 5x10^4^ vg) AAVs on DIV 7. A 488 *nm* excitation laser in combination with a 525*/*50 *nm* emission filter was used for EGFP imaging.

### Processing of extracellular data

The extracellular data from the long-term recording period and from the paired HD-MEA and whole-cell patch-clamp recording were spike sorted with SpyKING Circus (version=0.8.4; parameters: spatial radius considered=210 *µ*m, width of templates=3 ms, spike threshold=6, cut-off frequencies for band-pass Butterworth filter=300 Hz/9500 Hz). For long-term data, the 15 min recording chunks from each electrode set were concatenated (yielding a total recording time of at least 3 h) and separately spike sorted. The HD-MEA data of the paired recordings for synaptic input estimation, were separately spike sorted, followed by manual curation with the SpyKING Circus curation interface (using template similarity and spike time cross-correlogram characteristics as merging criteria). This procedure typically yielded clean units, based on which the synaptic input waveform estimation was performed. It was a robust approach, because even if some noise units with random spiking were retained, these units would likely be removed as there was no correlated postsynaptic activity. Moreover, if – in rare cases – two units had not been correctly merged, this would be revealed by input waveform estimates with very similar characteristics. Finally, the (presynaptic) units that were found to form a connections to the patched cell and the postsynaptic unit had to be identified in the long-term recording data. The HD-MEA footprint of the postsynaptic cell was generated by spike-triggered averaging (see Fig. 2D). For each of the presynaptic footprints and the postsynaptic footprint, we identified the corresponding best-matching footprints in the spike sorted long-term recording data (by using the maximum of the normalized cross-correlation between footprints). For almost all footprints, a clear match was found, and the unit was curated as above; only for very few individual footprints there was no match, and the respective connection was then excluded. An additional indication of the quality and validity of the spike sorting and footprint-matching procedures is given by the paired HD-MEA and IC whole-cell patch-clamp experiments (see Fig. S5). The extracellularly detected spike times of the identified postsynaptic unit matched the intracellularly recorded action potentials well, which held even true for high-activity periods with some variations in action-potential shapes.

### Regression approach for the estimation of synaptic input waveforms based on paired HD-MEA and patch-clamp recordings

We estimated synaptic input waveforms (e.g., EPSCs) by least squares linear regression of the whole-cell patch-clamp trace on simultaneously recorded unit spike trains according to equation 1 (see results section). Before running this estimation procedure, a preprocessing step to detrend the patch-clamp trace was applied. This was necessary as slow fluctuations appear as baseline shifts when short windows of the length of a single EPSC are concerned. These baseline shifts hinder the proper estimation of the waveforms. Briefly, the patch-clamp current trace was first down-sampled to 5 kHz and subsequently detrended. Detrending involved two steps. In the first step, very slow fluctuations in the current trace were determined using a sampling stride of 100 and applying a 5k-order median filter. The filtered trace captured very slow baseline drifts and was subsequently subtracted from the current trace. We call the modified current trace after this first detrending step *S_det_*_1_. For the second detrending step, periods of high synaptic activity in *S_det_*_1_ were detected and replaced by the median (*m̃*) value of the entire *S_det_*_1_ trace. To select high-activity periods, we calculated for *S_det_*_1_ the variability measure *ŝ* as *m.a.d.* (median absolute deviation) × 1.4826 and subsequently identified regions that were < *m̃*−3×*ŝ* or > *m̃* +3×*ŝ*. The remaining slow fluctuations were then determined by applying a 8k-order median filter to this modified *S_det_*_1_ trace. The resulting baseline trace was subtracted from *S_det_*_1_, which yielded the fully detrended current trace. The advantage of this two-step detrending procedure is that slow baseline fluctuations can be determined, even though the current trace features periods of high synaptic activity (e.g., due to network bursting). The prepossessing steps were applied to all VC whole-cell current traces shown here.

To verify that the preprocessing modifications to the current trace did not introduce major alterations in the EPSC waveform estimates, we developed an alternative approach for EPSC estimation based on spike-triggered averaging (STA) of isolated events (see **Fig. S8** and methods section below). The STA method identified drastically fewer connections, but did not require a modification of the current trace. The EPSC estimates of connections that were identified by both the regression and STA method matched well (**Fig. S8C**), suggesting that the effects of the preprocessing steps on waveform estimation were negligible.

A decisive advantage of our regression approach is that overlapping postsynaptic responses can be included in the EPSC waveform estimation. However, there are potential reasons (e.g., computational costs and non-linear interactions) to not include the most active periods of network spiking, which are associated with large postsynaptic currents. We evaluated the effect of different current thresholds, that determined which parts of the patch-clamp recordings were included in the regression analysis, and found 30 × *ŝ* of the current trace to be a good compromise, which was therefore used for all recordings (**Fig. S9**).

In the final step to estimate EPSCs, the spike trains of all units in the network with at least 10 spikes were encoded in a sparse binary u-by-t matrix (u: number of units; t: number of sampling points in the down-sampled current trace). Based on this spike time matrix and the preprocessed patch-clamp current trace, synaptic input waveforms were estimated with the *estimWaveforms* function of *Pillow et al. 2013* (https://github.com/pillowlab/BinaryPursuitSpikeSorting). The waveform estimates included a 20 ms baseline period before the presynaptic spike occurred. Units were accepted to form a monosynaptic connection with the patched cell when the absolute EPSC amplitude > 10 × s.d. of the baseline. Virtually all of these waveforms exhibited a typical EPSC shape (fast rising and slow decay phase; very few individual traces that clearly deviated from this shape were removed).

We chose here voltage-clamp recordings of synaptic currents as the basis for the synaptic input estimation in order to minimize effects of non-linear interactions and due to the fast EPSC kinetics. The input estimation, however, can also be performed based on current-clamp recordings of postsynaptic potentials (**Fig. S10**). For the estimation of EPSP waveforms, the procedure was the same as for EPSCs estimation except that only the first detrending step was performed (due to the relatively slow kinetics of synaptic potentials). Moreover, all recording periods with a voltage deviation up to 20 mV from baseline were included in the regression analysis, which excluded periods with postsynaptic action potential firing and strong synaptic activity.

Our MATLAB code that implements all preprocessing steps and the regression procedure is available at: https://github.com/neuroju/mea_patch_mapping.

### Validation of regression approach by simulation of ground-truth synaptic inputs

To simulate ground-truth synaptic inputs, we generated artificial presynaptic spike trains and accordingly added a defined EPSC waveform to our measured patch-clamp current traces. The defined EPSC was the mean waveform of the (onset-aligned) EPSCs from all monosynaptic connections that were identified for the given postsynaptic (patched) cell. The entire EPSC waveform was scaled to achieve a desired EPSC amplitude. The EPSC estimation performance is likely to be influenced by the time periods (e.g., periods of heightened vs. low network activity) during which the EPSCs occurred. Therefore, trains of simulated presynaptic spike times were generated in a semi-random manner as follows: First, for each paired HD-MEA and patch-clamp recording, the measured spike times from all units in the network were combined and binned (100 ms bin size). Each bin count was then divided by the total number of network-wide spikes, resulting in probabilities that a spike occurred in the respective bin. For each simulation of a spike time, one of these bins would then be semi-randomly selected according to the respective probabilities determined in the previous step. The precise simulated spike time was then a random time point within the selected bin. This procedure ensured that simulated spike trains followed the general profile of network-wide spiking activity.

For each paired HD-MEA and patch-clamp recording with artificially added ground-truth synaptic input, EPSCs were again estimated with our regression approach. The error between ground-truth EPSC and EPSC estimate was quantified as the mean deviation in percent. Specifically, ground-truth EPSC and EPSC estimate were first normalized by the respective amplitude of the simulated ground-truth EPSC. The mean deviation between these two waveforms within a 30 ms window starting at ground-truth EPSC onset specified the error. The mean error across all experiments (n=14) was eventually calculated for each parameter combination (number of simulated synaptic events and amplitude of simulated EPSC).

The *F*_1_ score was calculated as follows:

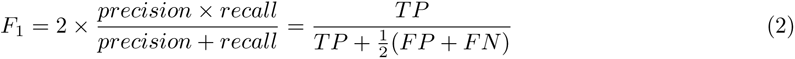

The number of true positives (TP) and false negatives (FN) was the number of accepted and excluded ground-truth connections, respectively (see previous section regarding acceptance criteria). To calculate the number of false positives (FP), we proceeded as follows: we added again artificial units with simulated spike times to all our paired HD-MEA and patch-clamp recordings as before, but this time simulated EPSCs were not added to the patch trace. The artificial unit had therefore no correlated activity in the patch trace. If the EPSC estimate of this simulated unit was nevertheless accepted by our regression approach, then it was counted as a false positive. The mean *F*_1_ score across all experiments was eventually calculated for each parameter combination.

With increasing EPSC amplitude and increasing number of synaptic events, the error between ground-truth and EPSC estimate quickly became marginal, and the F1 score reached values near its optimum of 1 (see Fig. 3D in results section). In purely experimental data, the number of synaptic events is dependent on the recording duration and the presynaptic spike rates. Our data set of paired recordings had a mean recording duration of 11.7 min *±* 7.0 min s.d., which – together with our simulation results – suggested that our approach would only have failed to identify connections with extremely small-amplitude EPSCs and very low presynaptic spike rates.

### STA approach for the estimation of synaptic input waveforms

Our alternative approach to estimate synaptic input waveforms, based on paired HD-MEA and patch-clamp recordings, relied on spike-triggered averaging of the patch-clamp current trace using only isolated presynaptic spikes (and the corresponding postsynaptic responses). By ’isolated’ we mean presynaptic spike times around which (20 ms before, 10 ms after) none of the other potentially connected units spiked. To obtain a sufficient number of events (*≥* 5) for averaging, unconnected units had to be identified and removed first: we iteratively calculated the STA EPSC for each unit (initially tolerating 256/128 spikes in the before/after window and then successively halving the number) and then removed the units that were within the spike tolerance when the absolute EPSC peak amplitude (*Amp_E_*) was smaller than 3 × s.d. of the pre-spike baseline and smaller than 5 pA. We repeated this procedure until no more units could be removed. Finally, STA EPSCs were calculated using only the isolated events. The remaining units were accepted as being connected to the patched cell when *Amp_E_ ≥* 3 × s.d. of the baseline and *Amp_E_ ≥* 5 *pA*.

### Reconstruction of synaptic conductance traces

In the main text, we described how the measured whole-cell current trace of the paired patch-clamp and HD-MEA and recording was reconstructed based on the EPSC estimates and corresponding presynaptic unit spike trains by applying the right term of equation 1. We can reconstruct the excitatory and inhibitory synaptic conductances experienced by a postsynaptic cell during the long-term HD-MEA recording of network-wide spiking – which preceded the patch-clamp experiments – in a similar way:

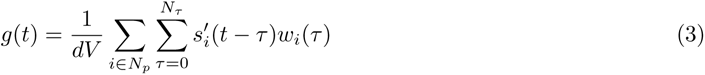

where *g*(*t*) is the synaptic conductance at time point *t*. As in equation 1, *w_i_* is the EPSC estimate for the i’th (presynaptic) neuron and *N_τ_* is the number of sample time points of the EPSC estimate; *s^′^* represents the corresponding (binary) presynaptic spike train. However, as opposed to *s_i_* from equation 1, which is the spike train of presynaptic neuron i from the paired recording, *s^′^* represents the spike train – of the same neuron i – from the long-term HD-MEA recording period. *dV* is the driving force (holding potential subtracted by the reversal potential of respective ion channels). *N_p_* specifies the set of presynaptic units that are included in the conductance trace reconstruction. For example, for the *g_i_* and *g_e_* trace reconstructions in Fig. 6, *N_p_* contains all inhibitory or excitatory presynaptic unit IDs, respectively, of a given postsynaptic cell. Similarly, the synaptic conductance of individual monosynaptic connections can be computed (e.g., for Fig. S4); in this case, *N_p_* contains only individual presynaptic unit IDs.

### Selection of high-conductance events

To calculate the mean conductance trace auto-(ACG) and cross-correlograms (CCG), we selected all high-conductance events that fulfilled the following criteria: both *g_e_* & *g_i_*were continuously for at least 300 ms > 2 × s.d. of the respective conductance trace.

To calculate the state-dependent spike-triggered averages of the inhibitory conductance trace, we defined *high g* states to be periods, during which *g_i_* > 3 × s.d. of *g_i_*trace, following smoothing with a 1k-point Gaussian filter. The remaining recordings period were considered *low g* states.

Similarly, periods with both high inhibition and high excitation were determined by *g_i_* > 2 × s.d. of *g_i_* trace & *g_e_* > 2 × s.d. of *g_e_* trace, using the smoothed g traces.

### Spike-transmission probability

The spike-transmission probability (STP) between two cells was always based on the long-term unit spike trains (> 3 h). For STP estimation, the cross-correlation between the spike trains of the two cells was calculated first (0.5 ms binning). The slow CCG ’baseline’ was determined by convolving the CCG with a partially hollow Gaussian kernel (standard deviaton=10 ms, hollow fraction=60%), as described before (English et al. 2017; Stark and Abeles 2009). Subsequently, this baseline was subtracted from the CCG. The baseline-subtracted CCG was then normalized with respect to the number of presynaptic spikes, and the STP estimate was the sum of bins during a certain window of positive lags. When the STP was extracted for the purpose of cell-type classification, a 4 ms window starting at the 1.5 ms positive lag was used.

To analyze the relationship between STP and connection strength, the exc/inhEPSC onset latency was used to determine the initial offset of the quantification window. Specifically, to account for differences in the response characteristics, a 4 ms window starting 0.5 ms before excEPSC onset latency was used for excitatory connections, and a 10 ms window starting 1 ms after the inhEPSC onset latency was used for inhibitory connections.

### Single-cell level data analysis of incoming connection properties

Several monosynaptic connection properties were determined. The spike rate of the presynaptic unit was the mean of the long-term recording period. The amplitude of the EPSC waveform estimate is the absolute difference between the EPSC peak and the waveform baseline. The EPSC onset latency was defined as the time difference between the time point at which the EPSC waveform reached a value below -5 × s.d. of the baseline and the presynaptic spike time.

### Extracellular unit footprint analysis

To characterize the extracellular electrical unit footprint of a neuron, i.e., the distribution of extracellular electrical potentials across the array electrodes, we generated the spike-triggered average for each of the simultaneously recorded traces (up to 1024; filtered at 0.3–9.5 kHz; spike cutout before/after was 5/10 ms) using the last 1500 spikes of the unit spike train from the long-term HD-MEA recording period. Such a number of spikes was recorded even from the neurons featuring the lowest spike rates. For each average electrode trace, we tested for negative peaks: we detected the first time point (*t*_0_), after spike time, at which the trace reached < -4 × s.d. of the trace baseline (first 2.5 ms); the peak was then the minimum value between *t*_0_ and *t*_0_+1 ms. Occasionally, especially for some excitatory cells, there were multiple negative peaks in the average electrode trace; possibly because the axonal signal was followed by the extracellular signal of an excited postsynaptic unit, or because the electrode recorded from multiple axonal branches. Our peak detection procedure ensured that only the first extracellular signal peak was used. An electrode trace was excluded if no peak was detected, and if there was no directly adjacent electrode with a detected signal. The signal latency was defined as the time difference between the signal peak of the respective electrode trace and the peak of the electrode trace with the largest (absolute) amplitude within the footprint, which typically was near the axon initial segment (AIS) and soma of the neuron (Bakkum et al. 2019).

AP propagation velocity was quantified based on the unit footprint as follows: for each electrode trace, we calculated the AP velocity at the given electrode position using the respective AP signal latency and using the distance – assuming a straight line – between the respective electrode and the cell soma (approximated by the electrode with the largest signal). We used the mean of these individual AP velocity measures to approximate the AP velocity of the neuron. Note that this AP velocity measure will be slightly larger than the true AP velocity, since we assume a straight axonal path between electrode and soma, while the true path will likely deviate from a line. Given this notion, our measure can be interpreted as capturing both AP propagation and how direct the axonal path is.

### Statistics

Analyses are based on data from 142 identified connections of 14 patched (excitatory) cells (1 inhibitory cell was excluded) from 5 culture preparations and animals. Statistical analyses were performed in R (R Core Team 2021) and MATLAB (MathWorks). Density scatter and violin plots in Fig. 6 were generated with https://github.com/aebergl/DensScat and https://github.com/bastibe/Violinplot-Matlab, respectively. Non-parametric two-tailed tests were conducted using the Mann-Whitney U test and Wilcoxon signed rank test. Student’s t-test was always conducted two-tailed. Normality was assessed with a Shapiro-Wilk test. Correlations were examined using Pearson’s R. The alpha level to determine significance was adjusted for multiple comparisons using the Holm correction when appropriate. No statistical methods were used to predetermine sample sizes. To probe the relationship of connection properties from the perspective of individual post-synaptic cells, we performed linear mixed-effects modeling using the R package *lme4* (Bates et al. 2015). Models contained spike rate and onset latency as as fixed effects with random slopes and intercepts, while postsynaptic cell ID was the random effect. A similar analysis was performed to examine the STP - EPSC amplitude relationship. Significance was assessed using a likelihood ratio test (Yu et al. 2021).

## Author Contributions

Conceptualization, investigation (unless otherwise stated), analysis (unless otherwise stated), writing of original draft: J.B.; Development of the regression method: F.F.; Intracellular-extracellular-spike analysis: S.S.K.; Confocal-microscopy investigations: A.P.B., X.X., J.B., K.C.K.; Assistance with patch-clamp experiments: T.G.; Spike-sorting support: M.S., T.K.; Supervision and funding acquisition: A.H.; All authors contributed to writing the final version of the manuscript. Additional analyses for the revised manuscript: J.B., S.S.K., T.G. The authors would also like to thank Vishalini Emmenegger and David Jäckel for their contributions in setting up the rigs for combined HD-MEA and patch-clamp recordings.

## Data availability

The data that support the findings of this study are available from the corresponding author upon reasonable request.

## Supplementary Materials

**Figure S1.**
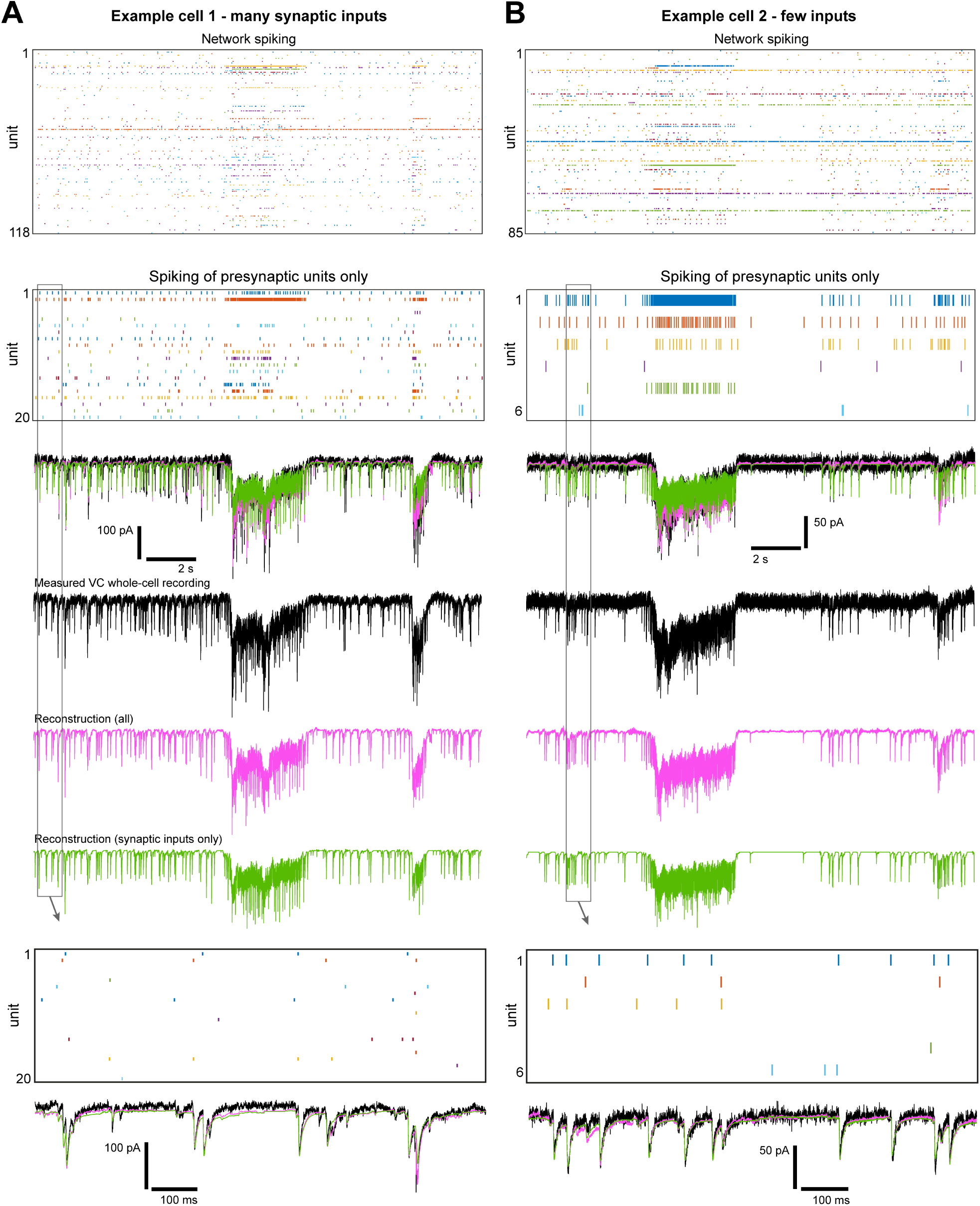
Variations in the number of identified synaptic inputs. Example synaptic input reconstructions of two neurons that were sequentially patched in the same network and yet displayed a difference in the number of identified incoming connections [**(A)** 20 vs **(B)** 6]. Example periods of unit spiking activity of the entire network and only of the respective presynaptic neurons (top two raster plots) are shown, in addition to the measured (black) and reconstructed (magenta/green) input-current traces. The current-trace reconstruction showed a good matching with the measured VC patch-clamp recording for both neurons, which suggests that the difference in identified inputs was of biological origin and not due to an incomplete input mapping.

**Figure S2.**
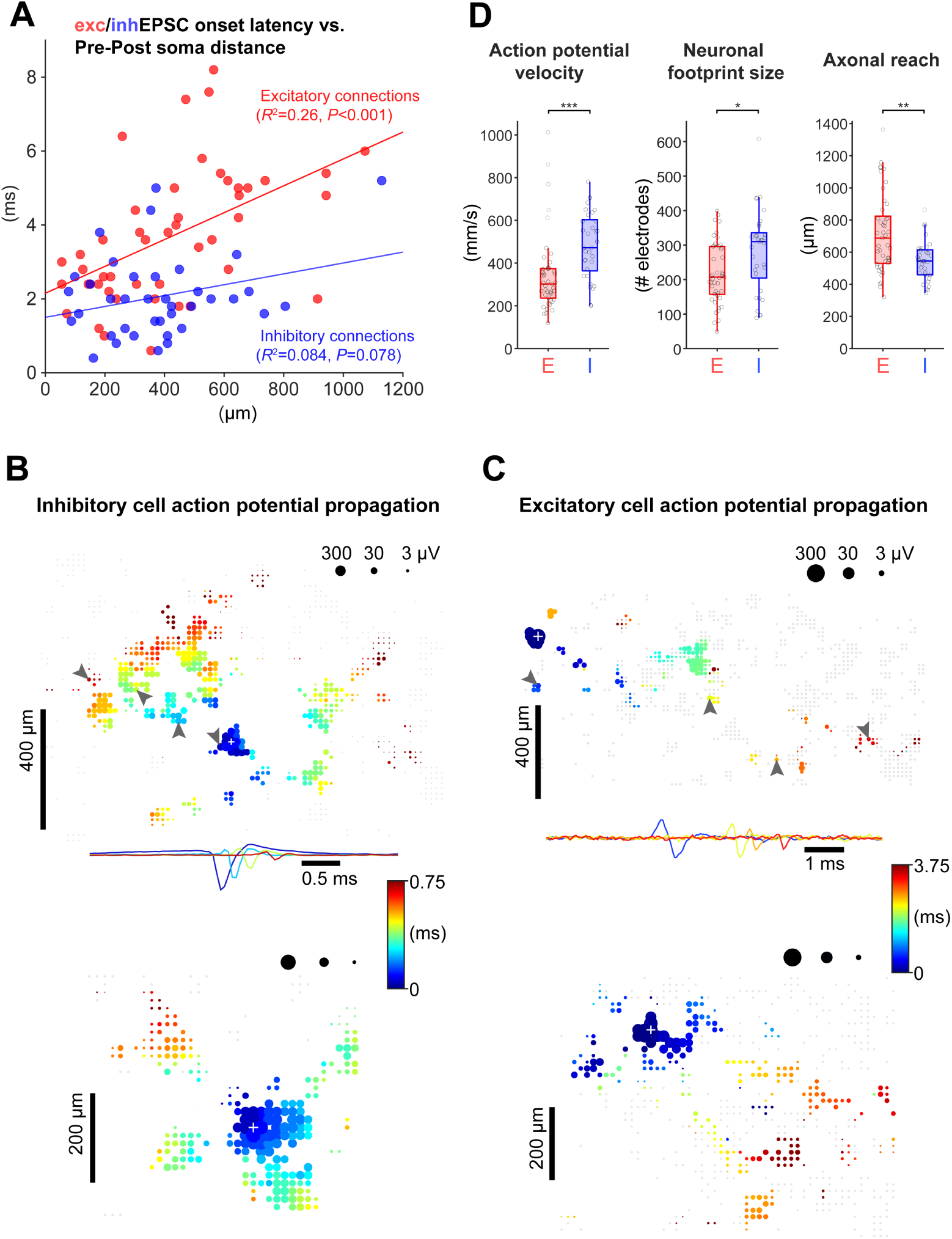
Inhibitory and excitatory neurons show differences in action potential propagation, neuronal footprint size and axonal reach. **(A)** Relationship between excEPSC (red) / inhEPSC (blue) onset latency (relative to presynaptic spike time) and the distance between pre- and postsynaptic cell. Lines represent linear regression fits ([E/I] 47/38 connections from 7 cells). **(B)** To visualize and quantify action potential (AP) propagation for a given unit, the spike-triggered average for all simultaneously recorded traces (up to 1024) was computed, and the peak latency relative to the electrode featuring the largest signal (marked by a white plus sign) was determined. Two example AP latency plots for inhibitory units with an extensive (top) and relatively small (bottom) axonal arbor are shown. Gray arrow heads in the top plot indicate the electrodes for which the STA traces are displayed underneath (filtered at 0.3–9.5 kHz). Gray dots indicate electrodes that did not show any AP signal. White ’gaps’ indicate electrodes that were not selected for long-term recording. **(C)** As in (B), but for excitatory units. **(D)** Difference between excitatory and inhibitory units in terms of action potential propagation velocity (*U* = 340, *P* < 0.001, Mann-Whitney U test), footprint size (*U* = 488.5, *P* = 0.010) and axonal reach (*U* = 425, *P* = 0.0013); [E/I] 45/33 pooled presynaptic units; for each unit, 1500 spiking events (# spikes available for all units) were averaged to generate the footprint. AP propagation velocity was quantified based on the AP latencies across the entire unit footprint (see Methods for details). Axonal reach is defined as the 90th percentile of the distances between the electrodes that belong to the unit footprint (i.e., show an AP signature) and the largest signal electrode (presumably near the soma). Box plots indicate median and interquartile ranges, and whiskers the minimum/maximum values except for outliers. \**P* < 0.05, \*\**P* < 0.01, \*\*\**P* < 0.001.

**Figure S3.**
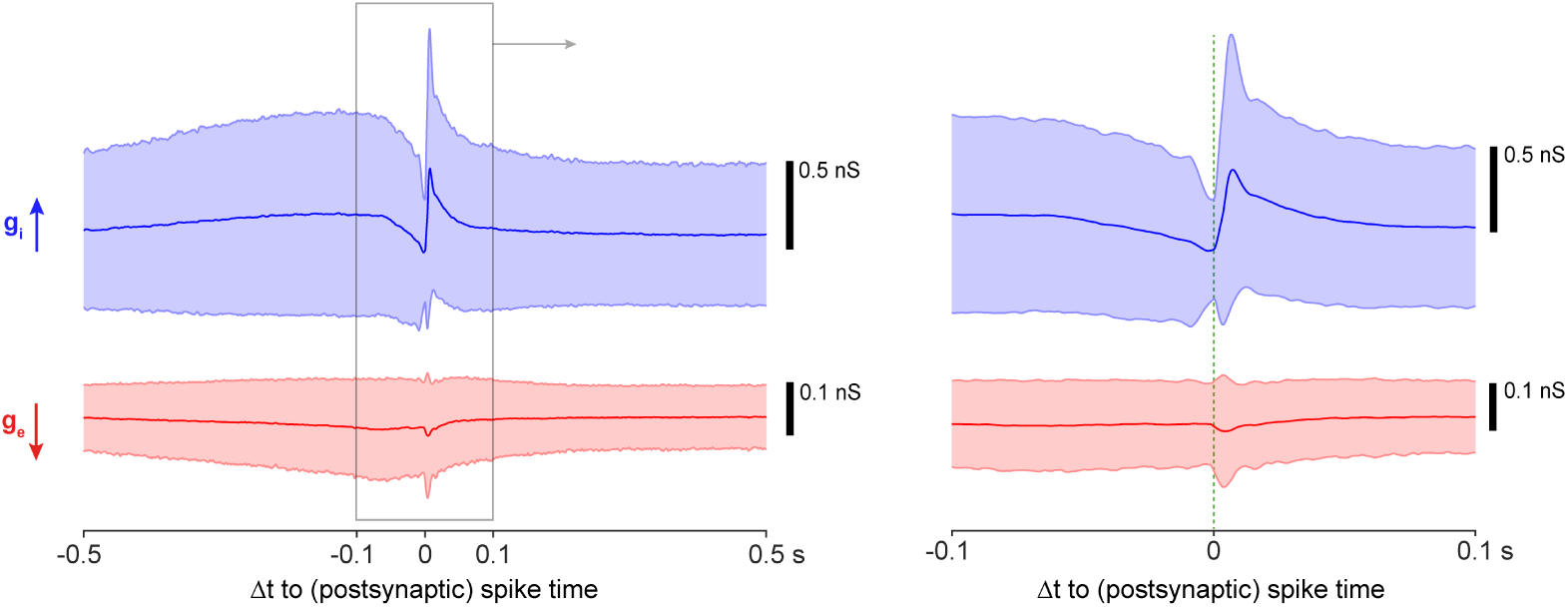
Variability in near-spike input conductances across entire recording. The spike-triggered-average conductances, *g_e/i,STA_*, from Fig. 6D shown with s.d. (shadings). Note that spike times from both low- and high-activity states were included (n = 15’813 postsynaptic spikes).

**Figure S4.**
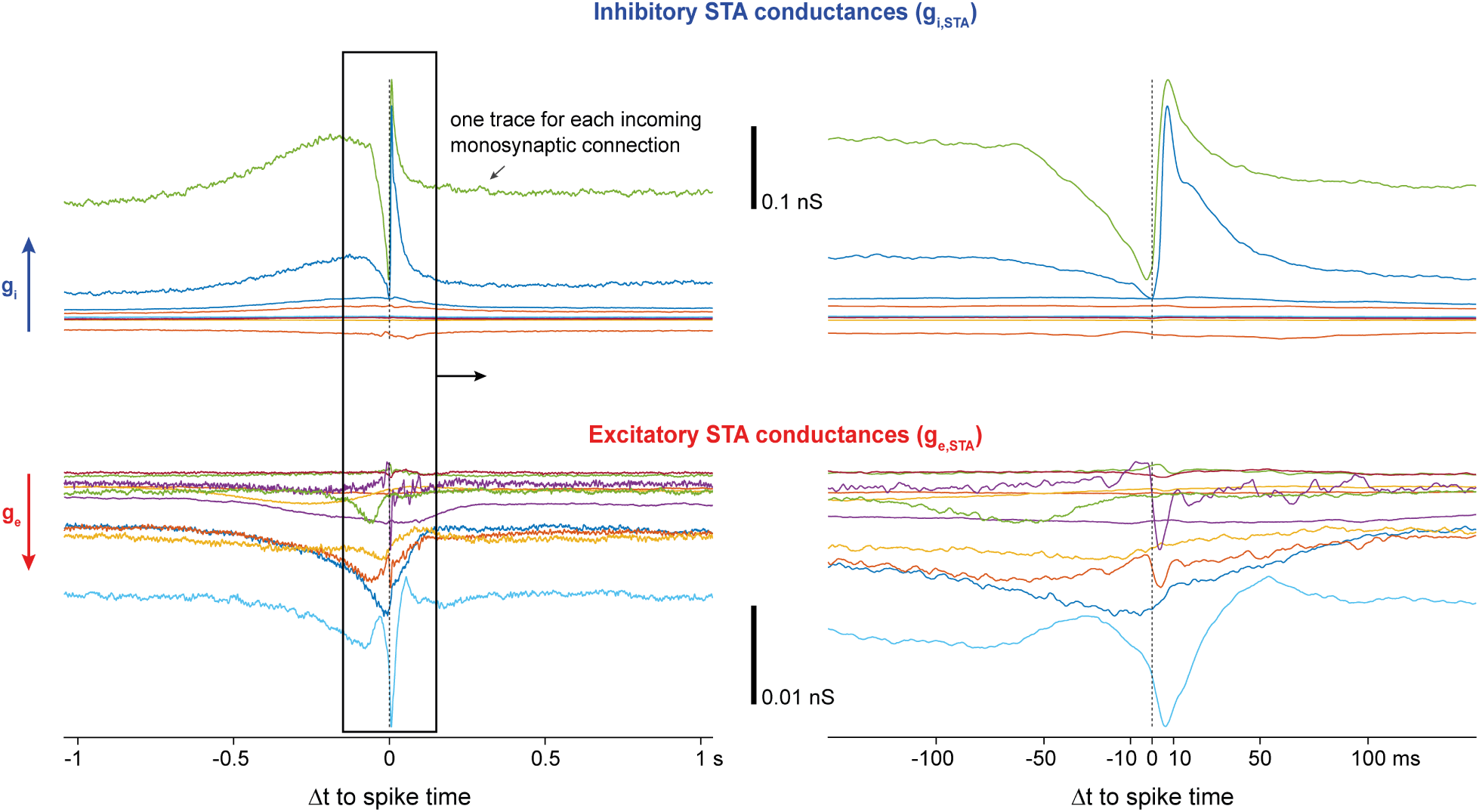
Role of individual incoming connections in controlling postsynaptic spiking. The spike-triggered-average conductance, *g_e/i,STA_*, was calculated separately for the conductance trace of each incoming connection to an example postsynaptic cell (9 inhibitory and 11 excitatory inputs). There appeared to be a few key inputs that had a particularly strong role in controlling postsynaptic spiking. Moreover, there was a subset of excitatory inputs with a sharp increase in conductance near postsynaptic spiking, while others only contributed through slow conductance changes. Interestingly, postsynaptic spiking typically occurred during the early rising phase of the rapid increases in excitatory conductance. As expected, the contribution of each connection to the total STA conductance depended on the respective synaptic strength and presynaptic spike rate: the mean *g_e/i,STA_* (from -2 s to +2 s) for each input correlated well with the respective product of input amplitude and presynaptic spike rate, for both excitation (*R*^2^ = 0.77; *P* < 0.001; n = 11) and inhibition (*R*^2^ = 0.96; *P* < 0.001; n = 9).

**Figure S5.**
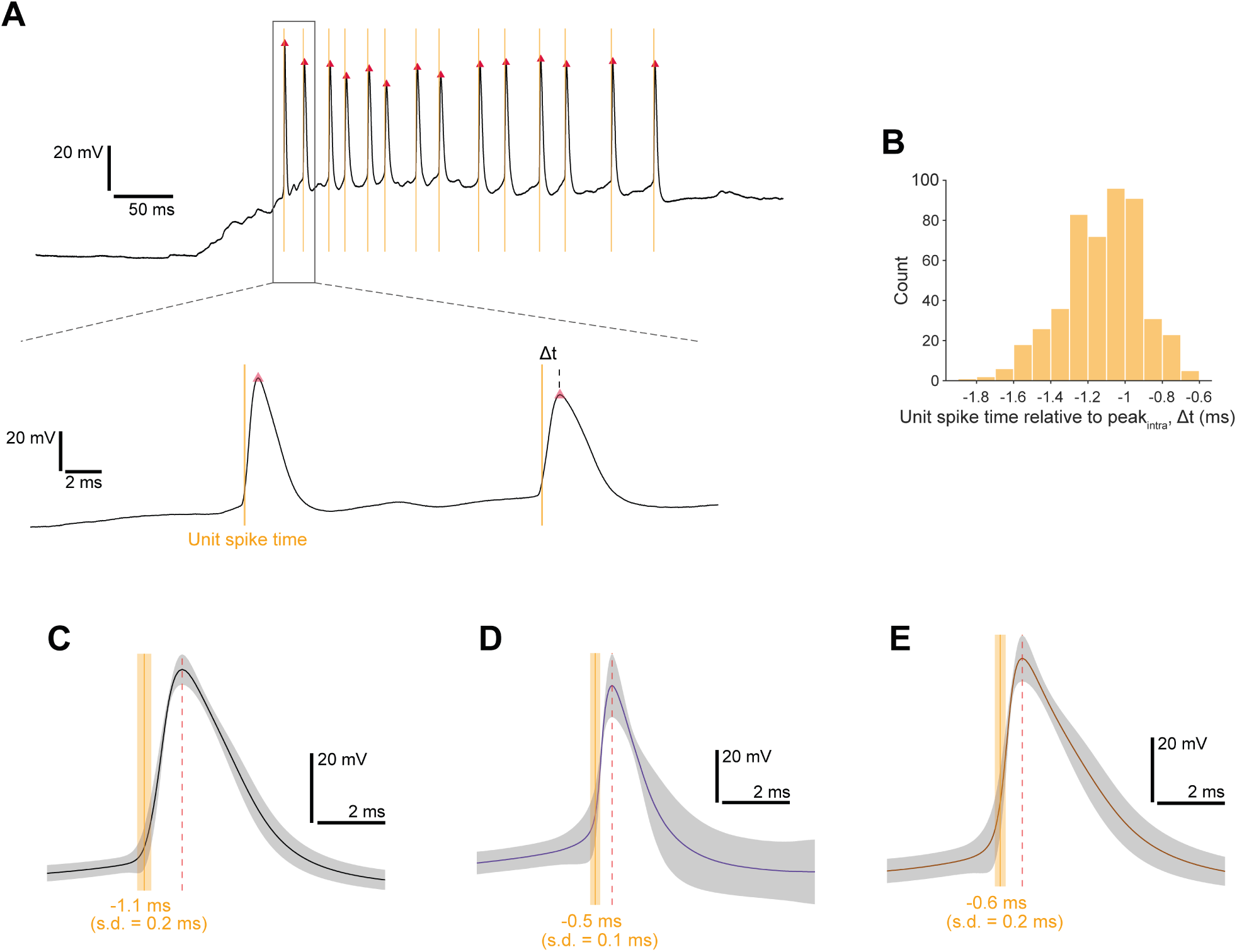
Relationship between extracellularly detected unit spike times and corresponding intracellular action potentials. Paired HD-MEA and IC whole-cell patch-clamp recordings during spontaneous spiking activity were performed, and the unit of the patched cell was identified by footprint matching (analogous to Fig. 2D), following spike sorting of the HD-MEA data. An IC internal solution with physiological chloride concentration was used. **(A)** Example patch-clamp voltage traces with action potentials. Red triangles mark the peak of the intracellular action potentials. Yellow lines indicate spike times of the unit that corresponded to the patched cell. **(B)** Histogram of the time differences between unit spike time and intracellular action potential peak of the same cell as in (A). **(C)** Mean action potential waveform and mean unit spike time (shadings: s.d.) of the same cell as in (A/B). **(D/E)** Results of two more cells, displayed as in (C).

**Figure S6.**
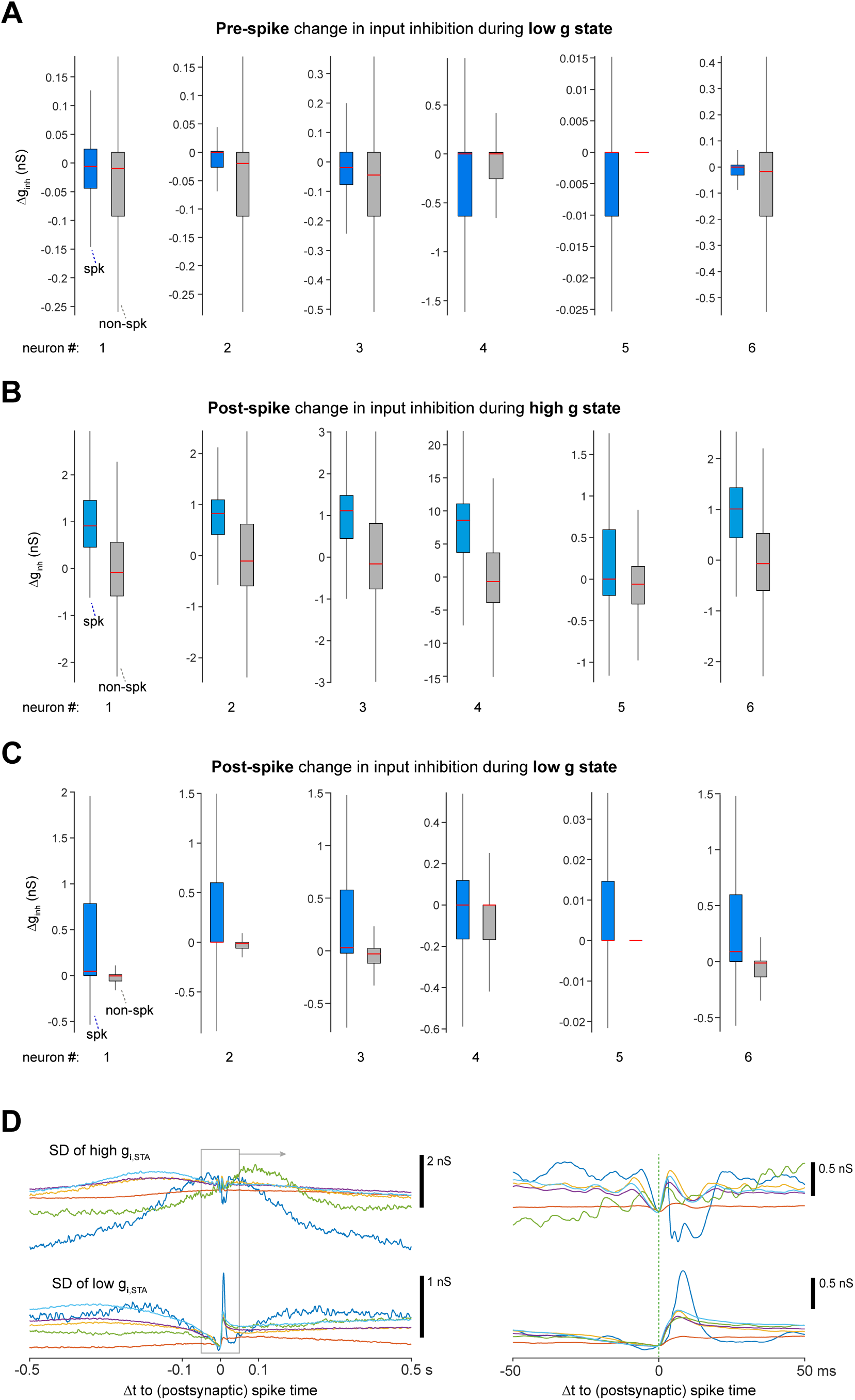
Overview of changes in input inhibition before and after individual postsynaptic spikes. **(A)** Analogous to Fig. 6J, the change in input inhibition before individual postsynaptic spikes was quantified for events that occurred during low-activity states. Often narrower ‘spk’ distributions could indicate restricted windows of opportunity for spiking (e.g., only after significant decay of any input inhibition). **(B/C)** Change in input inhibition right after postsynaptic spikes that occurred during high- (*B*) and low-activity (*C*) states. For this quantification, the mean *g_i_* value from t+5 ms to t+10 ms was calculated for each spike time t, followed by subtraction of the mean *g_i_* value from t-1 ms to t+1 ms. **(D)** Standard deviations across the spike-triggered average input inhibition traces, calculated separately for spiking events from high- (top) and low-activity (bottom) periods. Traces are aligned to the standard deviation value at spike time. Box plots indicate median and interquartile ranges and whiskers the minimum/maximum values except for outliers (outliers not shown). In the reference distribution of neuron 5 in (A/C), quartiles Q1-Q3 equal zero. By definition, a neuron experienced an inhibitory high-conductance event when *g_i_*> 3 × s.d. of *g_i_*. For non-spike reference distributions, time points were randomly selected from either high- or low-activity periods, excluding *±* 20 ms adjacent to measured spikes. It was n = 6 postsynaptic neurons in all panels.

**Figure S7.**
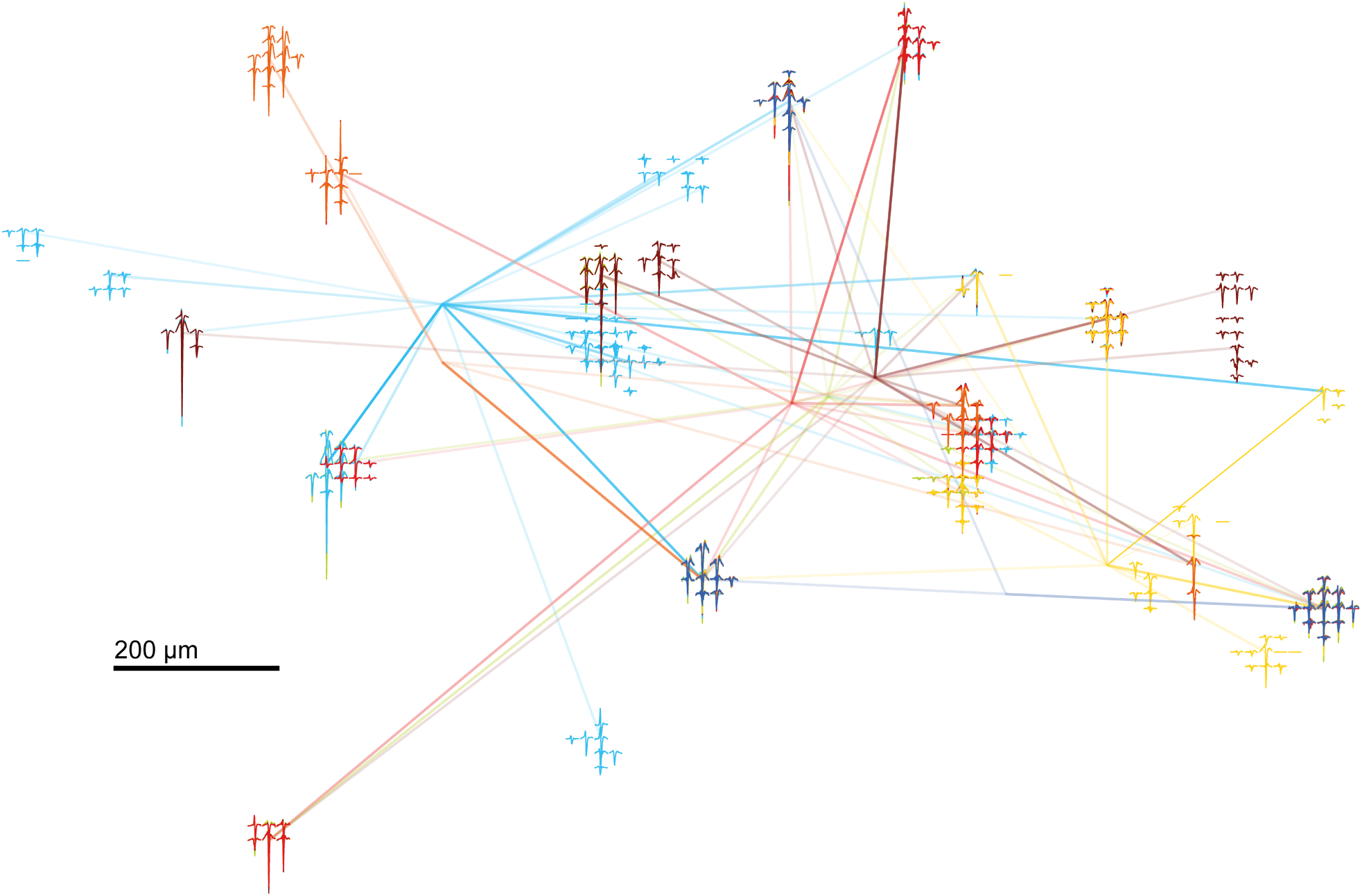
Spatial distribution of presynaptic unit footprints and monosynaptic connections that were identified based on multiple paired recordings in the same network. Equivalent to Fig. 9A, with unit footprints shown (filtered at 0.3–9.5 kHz). Only the traces from electrodes near the electrode with the largest signal amplitude are displayed for each unit. Note that some footprints match in terms of both location and signal characteristics, suggesting that the identified units represent the name neuron.

**Figure S8.**
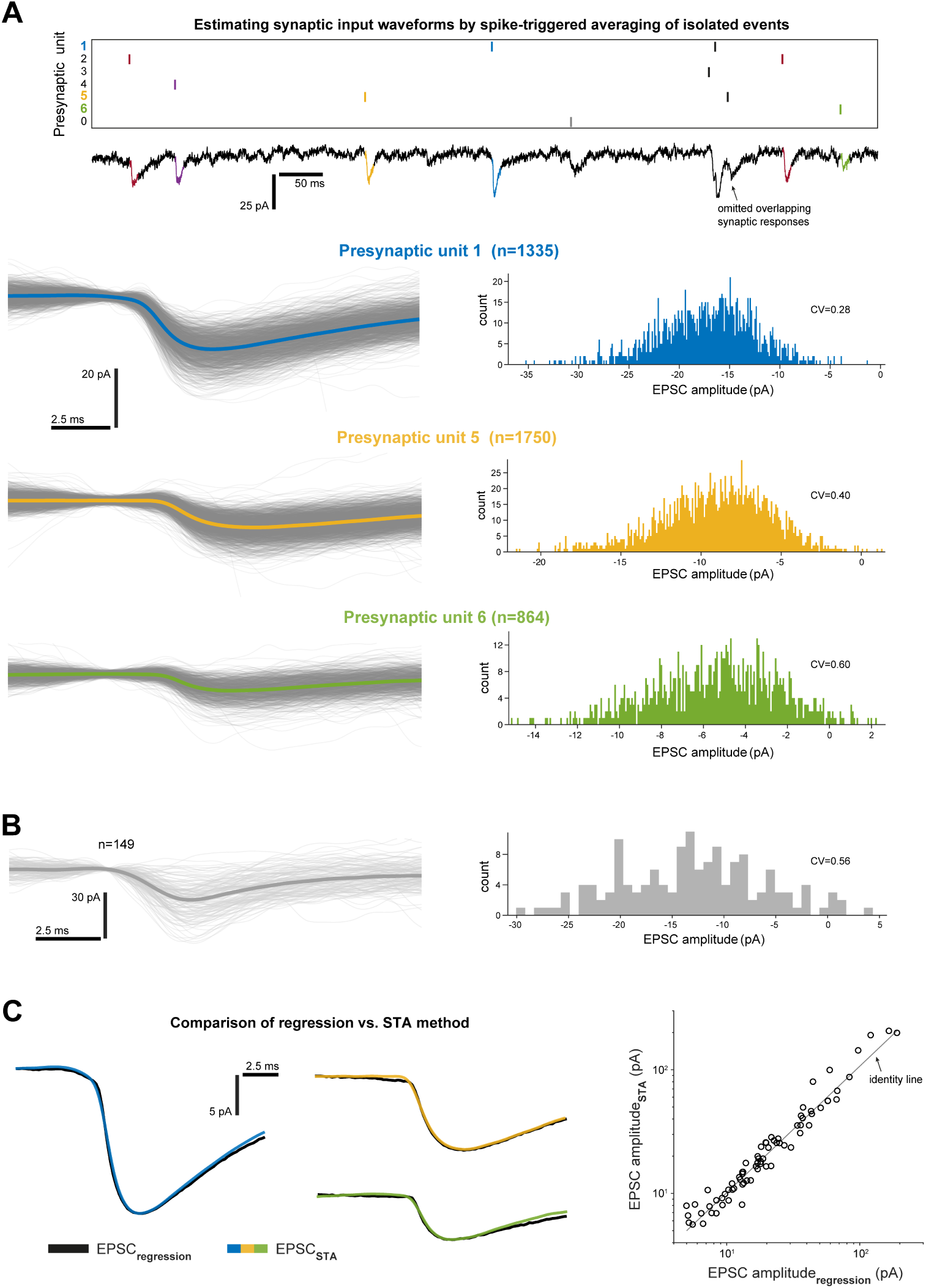
Spike-triggered-average-based estimation of EPSC waveforms using paired HD-MEA and patch-clamp recordings. An alternative to the regression-based EPSC estimation approach relied on calculating, for each unit in the network, the spike-triggered average patch response using only ‘isolated events’; that is, presynaptic spike times around which none of the other potentially connected units exhibited spiking (see Methods for details). **(A)** Top: example spike raster plot of presynaptic activity with corresponding whole-cell current-clamp recording. Isolated events are highlighted in colour. Overlapping responses (arrow) were excluded. (‘Unit 0’ contains spike times from units that might be connected to the patched cell, but without reasonable confidence). Bottom: for three example inputs, the mean (colored) and the individual EPSCs (gray) across the entire recording are shown, in addition to the respective EPSC amplitude histogram (a Gaussian filter with a 20-element sliding window was applied to individual response traces for better visualization; mean trace based on raw data). **(B)** Another example of evoked EPSCs and amplitude histogram from a separate recording. ‘Peaks’ in the amplitude distribution consistent with multi-site quantal release. **(C)** Left: Comparison of EPSC-waveform estimates based on regression vs. STA method of example connections shown in (A). Right: EPSC amplitudes, based on connections identified by both methods, were highly correlated.

**Figure S9.**
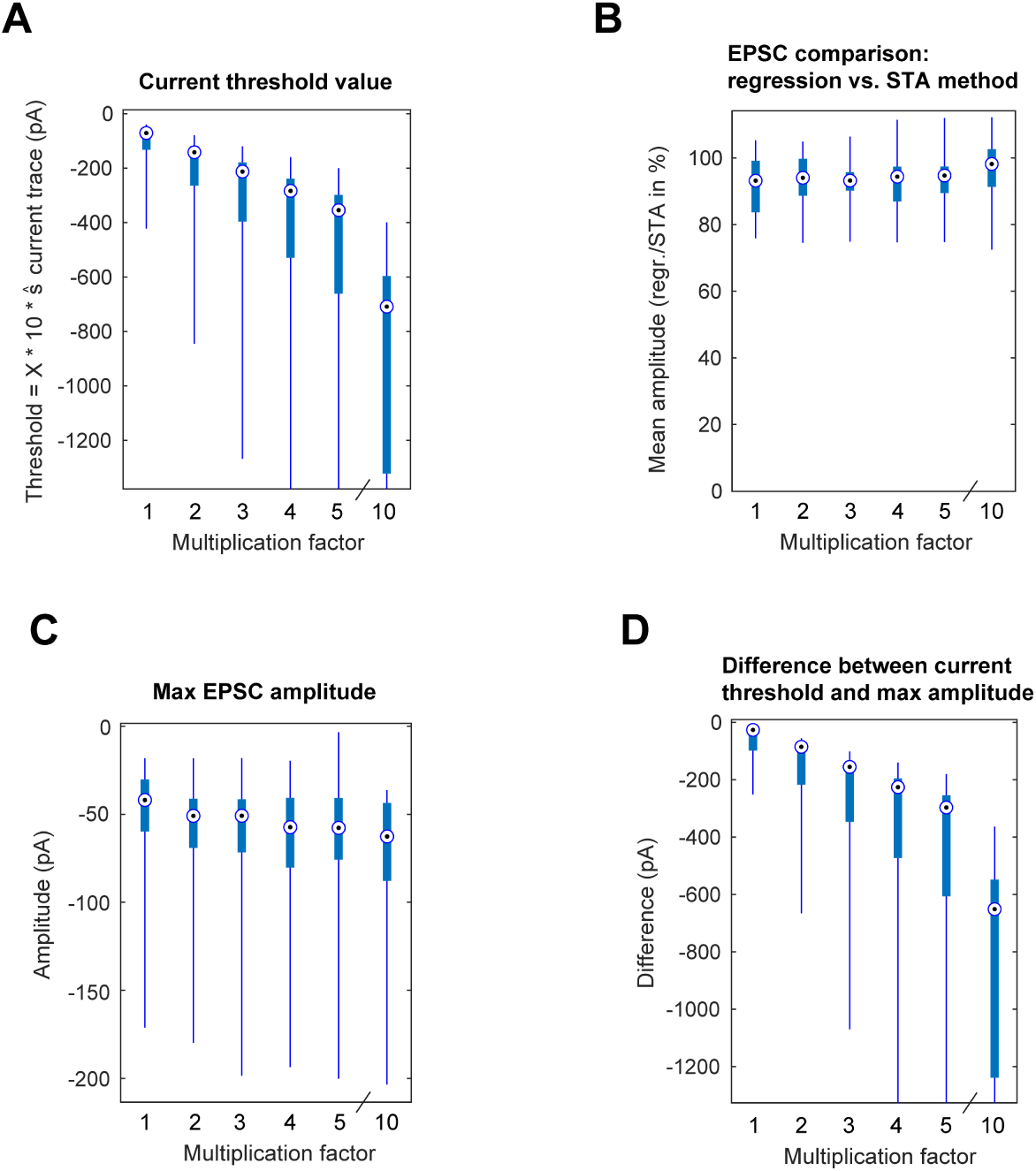
Effect of inclusion current thresholds on the estimation of EPSC waveforms. Different current thresholds, that determined which parts of the recording were included in the regression analysis, were applied to the patch-clamp current traces. The threshold values were multiples of the standard deviation of the respective current trace. Specifically, for each recording, 6 threshold values were evaluated: 1, 2, 3, 4, 5 & 10 × 10 *× ŝ* of the current trace. With *ŝ* as *m.a.d. ×* 1.4826. **(A)** The determined current thresholds across experiments. **(B)** We used the EPSCs determined by our STA method, which was performed without any inclusion criteria involving current thresholds, as ground-truth EPSCs. To these ground-truth EPSCs we compared the regression-EPSC estimates that were determined using the different current thresholds. Specifically, for each paired recording, we calculated for each common synaptic input the ratio of the EPSC amplitudes determined by the regression and STA method. The mean of these ratios was calculated for each postsynaptic cell, which is summarized here. **(C)** Largest EPSC amplitude for each cell. **(D)** Difference between current threshold and largest EPSC amplitude. In all panels, box plots indicate median and interquartile ranges, and whiskers the minimum/maximum values (no outliers). Some extreme whiskers in (A) and (D) were trimmed for better data visibility. [14 cells in (A-D); 9 patched cells in (B), where recordings were only included if at least three common inputs were identified by using the two methods.] Compared to the EPSC amplitudes determined by the STA method, the different current thresholds had only a small effect on the EPSC amplitudes that were determined by the regression-based method (B). Moreover, the maximum EPSC amplitude that was identified showed little dependence on the current threshold (C). These findings suggested that synaptic inputs exhibited sufficient activity – i.e., for a reliable EPSC estimation – during periods that were selected by even the smallest current thresholds. Moreover, the current threshold values were well above the maximum EPSC amplitudes as confirmed in (D). We chose a threshold of 30 *× ŝ* of the current trace, which resulted in a median current threshold of approximately 200 pA, with sufficient distance to the maximum EPSC amplitude, while also keeping computational costs within a feasible range.

**Figure S10.**
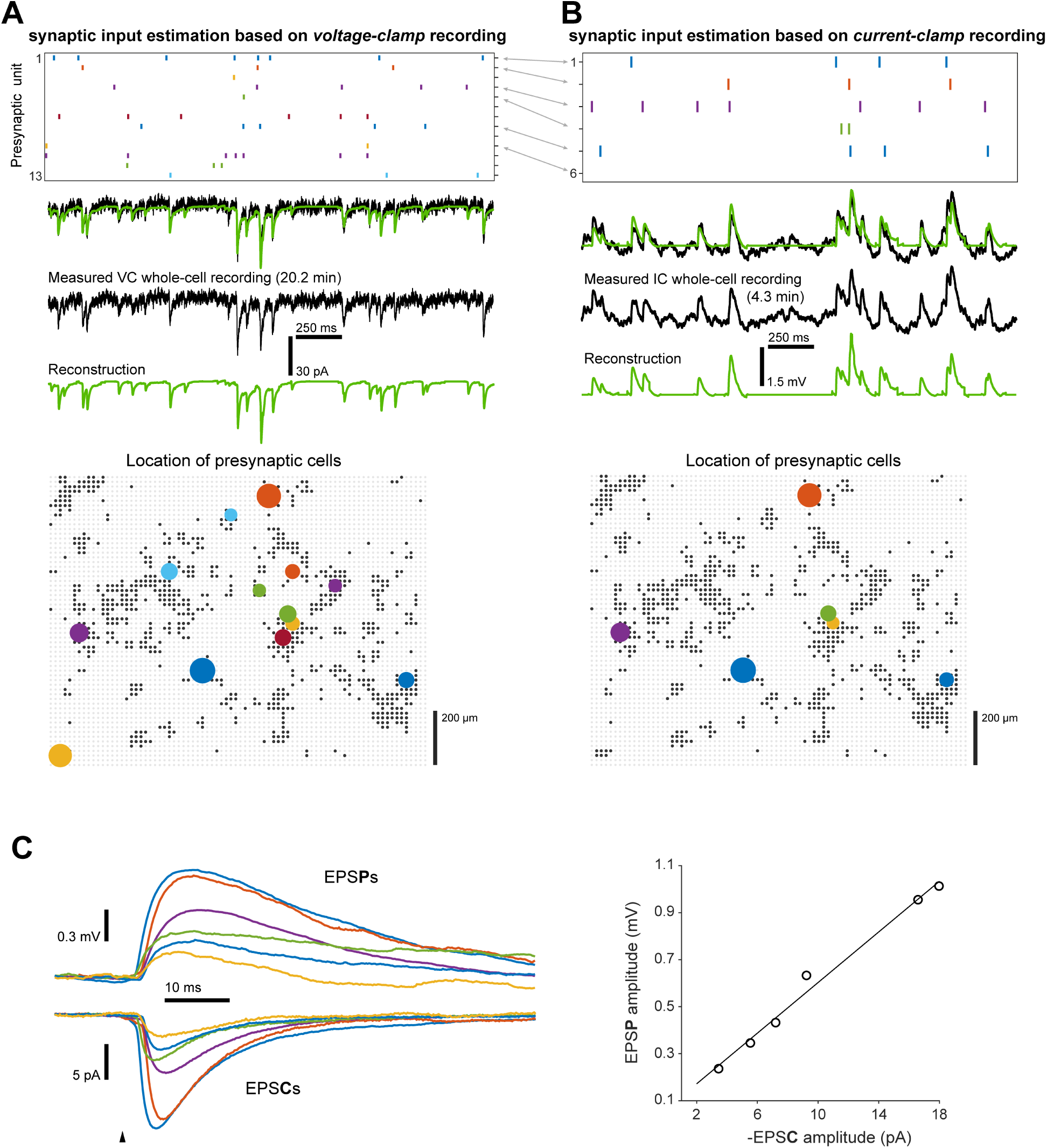
Estimation of synaptic input waveforms based on current-clamp recordings. Two paired HD-MEA and whole-cell patch-clamp recordings were acquired from the same cell in either voltage-clamp **(A)** or current-clamp mode **(B)**. **(A/B)** Top: raster plot of the identified presynaptic units of an example period (corresponding presynaptic cells are marked by grey arrows). Middle: patch-clamp recording (black) and reconstruction based on synaptic inputs (green). Bottom: location of presynaptic cells; one coloured filled circle per cell (black dots: HD-MEA electrodes selected for recording; grey dots: electrodes not recorded from). The same presynaptic cells identified in (A) and (B) have matching colours. The larger the circle the stronger the connection. **(C)** Left: Corresponding EPSP- and EPSC-waveform estimates (waveforms from the same input have matching colors). Right: Correlation of EPSP and EPSC amplitudes (linear regression fit; *R*^2^ = 0.99).

